# Single-molecule studies of conformational states and dynamics in the ABC importer OpuA

**DOI:** 10.1101/2020.08.07.241463

**Authors:** Konstantinos Tassis, Ruslan Vietrov, Matthijs de Koning, Marijn de Boer, Giorgos Gouridis, Thorben Cordes

## Abstract

The current model of active transport via ABC importers is mostly based on structural, biochemical and genetic data. We here establish single-molecule Förster-resonance energy transfer (smFRET) assays to monitor the conformational states and heterogeneity of the type-I ABC importer OpuA from *Lactococcus lactis.* We present data probing both intradomain distances that elucidate conformational changes within the substrate-binding domain (SBD) OpuAC, and interdomain distances between SBDs or transmembrane domains. Using the methodology, we studied ligand-binding mechanisms as well as ATP and glycine betaine dependences of conformational changes. Our study expands the scope of smFRET investigations towards a class of so far unstudied ABC importers, and paves the way for a full understanding of their transport cycle in the future.

## Introduction

ATP-binding cassette (ABC) transporters represent the most abundant and diverse family of transport proteins known. They play crucial roles in numerous cellular processes including nutrient uptake[1], antibiotic and drug resistance[2], antigen presentation[3], cell-volume regulation[4] and others[5–9]. Despite their importance, the majority of molecular models proposed for transport are based on the functional interpretation of static crystal structures, due to the inability of classical biophysical and biochemical techniques to visualize dynamic structural changes[5–9]. Advanced mechanistic insights, for instance knowledge on structural dynamics and heterogeneity of conformational states in transporters, could be beneficial for the fight against pathogenic bacteria [10], and for the treatment of ABC related diseases such as cystic fibrosis[11] and multi-drug resistance in cancer cells[2]. Determining the structural dynamics of drug targets would allow the rational design of high-affinity drugs, and provide a better understanding of a drug’s mode of action[12]. However, the development of methods to make a membrane-embedded transport system amenable to complex biophysical investigation represents a huge bottleneck.

Over the past years[13], various labs have introduced single-molecule tools to investigate the structural dynamics of active membrane transporters[14–25]. Förster resonance energy transfer in combination with single-molecule detection (smFRET[26–28]) has proven to be a particularly useful tool for the validation of structural models[29–31], and for revealing functional features of transporters which are mechanistically important, such as conformational heterogeneity[28, 32–34]. While the labs of Shimon Weiss[14], Scott Blanchard[15–17] and Antoine van Oijen[18] have published the first smFRET studies of secondary active transporters in a detergent environment[14–16] and within liposomes[17, 18], we introduced smFRET for studies of ABC transporters in collaboration with the groups of Bert Poolman, Konstantinos Beis and Robert Tampé. These transporters included importers (GlnPQ[19, 20]) and their substrate binding proteins and domains[21] and the exporter McjD[23]. In addition, we have also studied the non-transporting ABC-protein ABCE1[35]. Lewinsson[24] and Slotboom[36] have performed detailed smFRET studies on the type-II ABC importer BtuCD-F. And the influence of different membrane mimics (detergent, lipid nanodiscs and proteoliposomes) was presented in a recent study on the ABC exporter MsbA[25]. Advances in the visualization of structural dynamics in transporters have also been made possible by other biophysical techniques such as highspeed AFM[37] and EPR[38, 39].

All these studies were motivated by the desire to answer long-standing key questions in the transporter field. These questions include ligand binding mechanisms, the mechanistic basis of substrate selectivity, the precise timescales of conformational changes in transmembrane and nucleotide binding domains, and the comparative specific conformational states at rest for the distinct type-I and type-II ABC importers. Furthermore, it remains a goal of researchers in the field to establish complete models of transport that reflect and enhance an understanding of the coordination of transport, ATPase activity, and the associated conformational changes. Such models would provide a precise understanding of how substrate binding and ATP hydrolysis events are coupled and transmitted from one domain to another via conformational changes that ultimately drive substrate transport.

We here extend our smFRET work on ABC-transporters to the osmoregulatory OpuA. This importer represents a well-established model system for the type-I ABC transporters and is involved in cell volume regulation via the uptake of glycine betaine [4, 40]. The domain organization of the transporter is shown in Figure 1a. Poolman and co-workers showed that OpuA is more complex than other type-I ABC importers such as the molybdate and maltose permeases because of the presence of two additional cytosolic domains (Cystathionine β-Synthase; CBS)[4, 40]. Each homodimer of OpuA consists of two subunits: one composed of the transmembrane domain TMD and substrate binding domain SBD (OpuABC), and the other OpuAA of the nucleotide binding domain (NBD) and CBS in tandem (Figure 1a). The CBS domains play a role in sensing intracellular ionic strength and in conjunction with an anionic membrane surface, gate the transporter in response to osmotic stress for import of glycine betaine. The cycling of the NBDs from a dimeric ATP-bound state to a monomeric ADP-state is probably dependent on the CBS-CBS interaction, which is regulated by ionic strength and in particular potassium[4, 40, 41]. The presence of the additional CBS domains might lead to a different mechanism of transport and/or transporter activation than occurs in other type-I ABC transporters. OpuA is a paradigm for osmoregulatory ABC transporters and the system is widespread in the bacterial and archaeal kingdoms[4]. It is critical for the survival of low GC Gram-positive pathogens under conditions of osmotic stress, but also plays an important role in osmoregulation in Gram-negative pathogenic bacteria[4].

**Figure 1.**
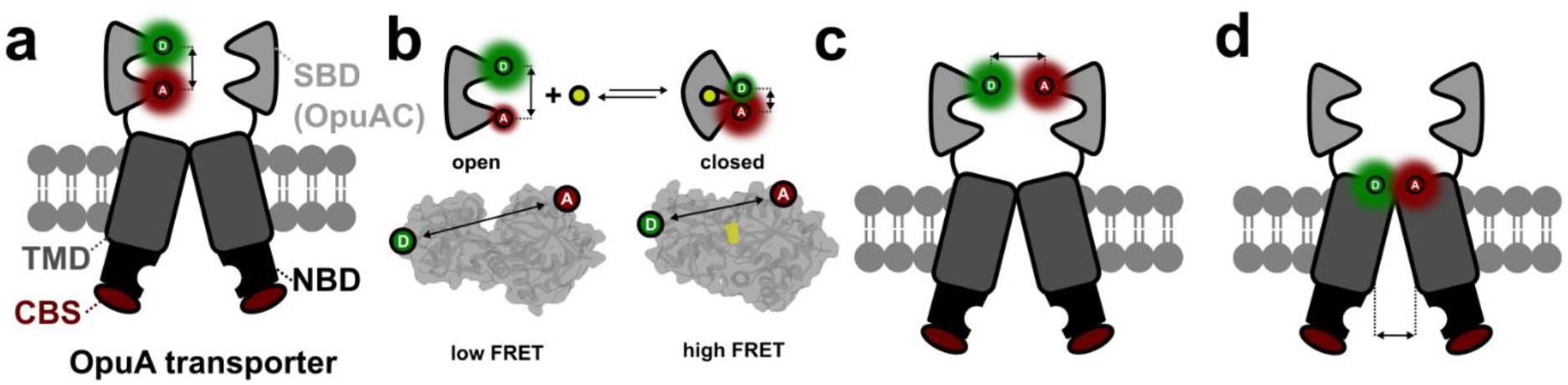
Setup for smFRET studies of the type-I ABC importer OpuA. **(a)** Structural organization of OpuA and intradomain FRET assay with the SBD. Please note that in our work, the second SBD can also be labelled, but this is not shown for clarity. **(b)** FRET assay principle to probe the conformational states of the SBD OpuAC via FRET. Open and closed structures are based on the published PBD structures of soluble OpuAC 3l6g and 3l6h, respectively, showing cysteine variant OpuAC (367C/423C) and ligand in yellow. **(c/d)** Interdomain FRET assays based on **(c)** SBD and **(d)** TMD labelling. Reactive coordinates probed in each panel are indicated by arrows and dashed lines.

OpuA is a suitable model system for the present work because it can be functionally reconstituted in nanodiscs, where it shows ion- and substrate-dependent ATPase activity with a high coupling efficiency, when nanodiscs are prepared with physiological lipids[40]. In this study, our aims were to monitor by confocal smFRET in solution [19–21, 23] various aspects of OpuA conformational dynamics (Figure 1) and to examine whether any structural heterogeneity exists. We examined the conformational dynamics of the SBD OpuAC in isolation, and in the context of the full transporter, and found that its ligand-binding behaviour was similar (Figure 1a/b), further supporting the utility of studies on SBDs and SBPs in isolation (as done previously for various ABC-related SBPs/SBDs[21]). OpuAC showed rapid ligand release times, a feature that might be argued to facilitate high transport rates[40, 42–45]. We also studied interdomain interactions between SBDs (Figure 1c) and evaluated suitable labelling positions for smFRET of the TMDs (Figure 1d). SBD-SBD interactions were examined in the ligand-free, ATP-bound, and glycine-betaine-bound states. Finally, for TMD studies, we relied on homology modelling of the TMDs aiming to label the extremes of transmembrane helices on either side of the membrane and generated a large-number of cysteine derivatives. These were screened to identify positions with high labelling efficiency and useful biochemical activity. We provide preliminary smFRET data on the most promising OpuA TMD mutant, from which no mechanistic conclusions could be drawn yet.

## RESULTS

To determine whether isolated OpuAC retains native ligand binding behaviour, we compared the isolated SBD (Figure 1b) with the SBD present in full-length OpuA (Figure 1a). To this end, we generated two double cysteine derivatives of OpuAC: OpuAC (360C-423C) also used in ref. [21] and OpuAC (367C-423C). Mutations were located on the rigid subdomains of OpuAC and substrate binding was expected to increase the proximity of donor and acceptor fluorophores on labelled OpuAC, i.e., lead to a high FRET efficiency state upon ligand binding (Figure 1b). Our goal initially was to maximize the FRET-efficiency change during the conformational motion, and so the impact of mutations or fluorophore labelling on OpuAC’s ability to dock with the transporter was not considered.

### Conformational dynamics of isolated OpuAC

To characterize the conformational dynamics of soluble OpuAC labelled with Cy3b and ATTO647N, we used both alternating laser excitation (ALEX) on diffusing molecules and confocal scanning microscopy on surface-immobilized ones[21]. In agreement with previously published results[21], OpuAC (360C-423C) changed its conformation in a ligand-dependent fashion (Figure. 2a-c).

**Figure 2.**
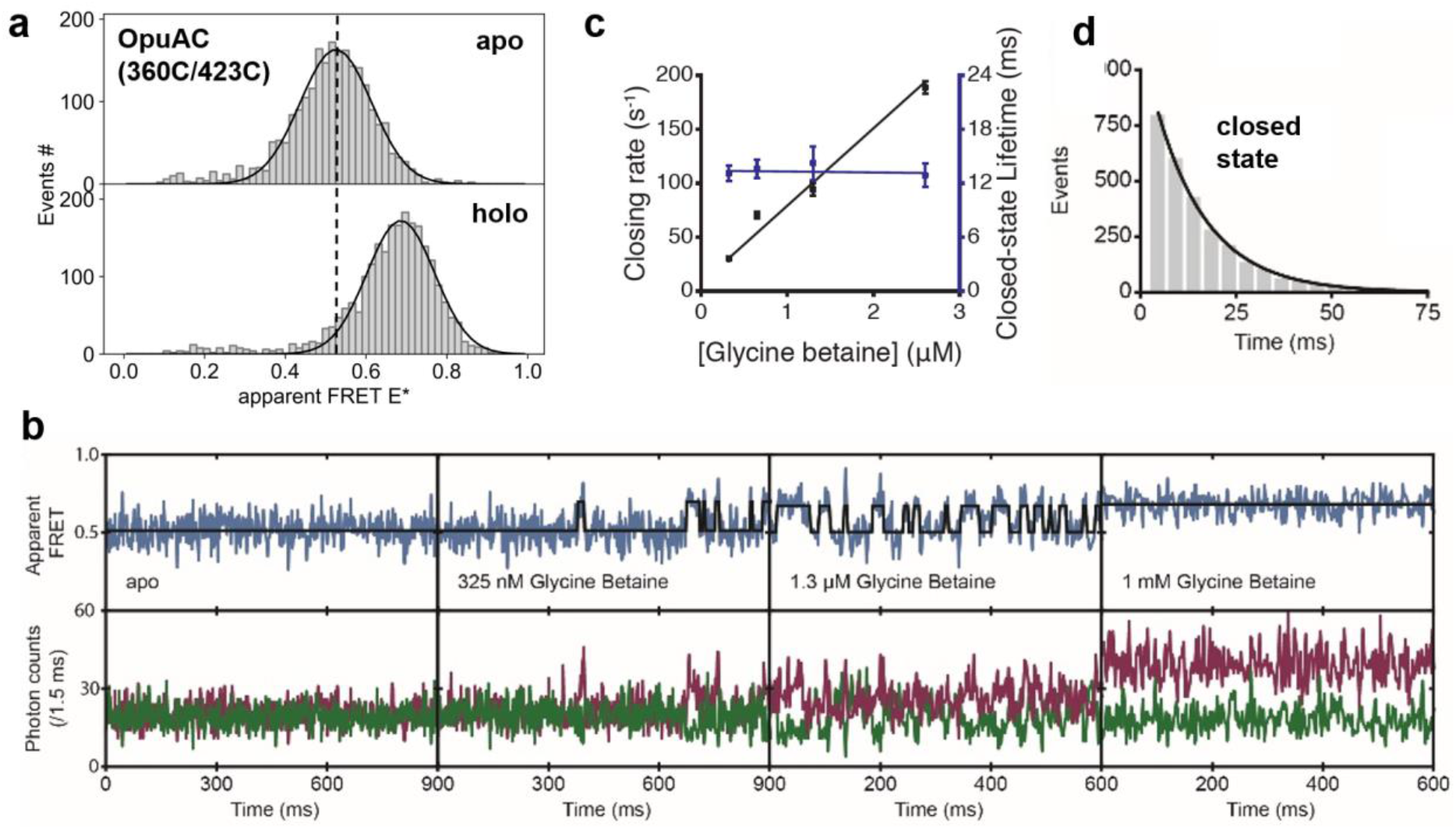
smFRET studies of isolated OpuAC (360C-423C) with Cy3B and ATTO647N as donor and acceptor fluorophores, respectively. **(a)** Apparent FRET efficiency histogram of freely diffusing fluorophore-labelled OpuAC molecules, obtained from the solution-based smFRET and ALEX measurements under the indicated conditions. **(b)** Fluorescence trajectories of OpuAC under different conditions as indicated; donor (green) and acceptor (red) photon counts were binned with 1.5 ms. The top panel shows calculated apparent FRET efficiency (blue) with the most probable state-trajectory of Hidden Markov Model (HMM) (black). **(c)** Average closing rate (black) and lifetime of the closed-state (purple) as function of glycine betaine concentration. Error bars indicate the 95% confidence interval. **(d)** Lifetime distribution of the closed state as obtained from the most probable state-trajectory of the HMM of all molecules. Grey bars are the binned data and the solid line is an exponential fit.

Both the open-unliganded (low FRET) and closed-liganded states (high FRET) were detectable and their occupancy changed as a function of the glycine betaine concentration in the buffer solution (Figure 2a/b). In the absence of glycine betaine, we find exclusively occupation of the open state (Figure 2a/b – apo), as was also shown previously[21]. This result and its interpretation were valid for freely diffusing (Figure 2a) and surface-immobilized OpuAC (Figure 2b). Identical results were obtained when an alternative pair of fluorophores was used for labelling (Alexa555/Alexa647, Figure S1). This suggests that the fluorophores do not compromise the conformational changes of OpuAC and do not alter the ligand binding affinity. The supporting data in Figure S1 also shows the two-dimensional character of the ALEX experiments, i.e., low stoichiometry acceptor-only molecules (S < 0.3), intermediate stoichiometry donor-acceptor-labelled molecules (0.3 < S < 0.8) and high stoichiometry donor-only molecules (S > 0.8). For histograms shown in Figure 2a and subsequent figures where 1D-E* histograms are presented, we focussed on the analysis of the FRET-efficiency distributions in the intermediate stoichiometry region, where the protein carries one donor- and one acceptor fluorophore.

Next, we studied the dynamic conversion between conformational states (Figure 2c). For this, the temporal evolution of donor and acceptor fluorescence signals of surface-immobilized (as described in the surface microscopy section in methods) OpuAC molecules were followed using confocal scanning microscopy. The FRET efficiency time traces show frequent switching between the low and high FRET state, which indicates the opening and closing transitions of OpuAC (Figure 2b). To obtain the associated kinetics, the time traces were fitted with a two-state hidden Markov model (HMM). The lifetime distributions of the low and high FRET states were obtained from the fit (as described in ref. [21], Figure 2c/d). From this, we extracted the average opening and closing rates (Figure 2c/d). We observed that the average opening rate was largely concentration independent, whereas the average closing rate scales approximately linearly with glycine betaine concentration. This ligand dependency is indicative of an induced-fit ligand-binding mechanism[19–21, 23], yet the limited time-resolution of 1.5 ms cannot rule out faster conformational switching under the conditions tested. We stress, however, that the fluorescent time traces of OpuAC (360C/423C) are of very good quality and OpuAC has the fastest ligand-release time that we have detected with HHM-analysis and dwell-time analysis amongst all SBPs and SBDs that we have analysed [21]. Our data (see below) also suggest that the ligand-binding affinity (K_d_) was in the low micromolar range, (Figure 2c), as determined by titration experiments on freely diffusing molecules. This ligand affinity was in agreement with a value from published bulk experiments where a K_d_-value of ~4 μM was reported[46].

### Conformational states and dynamics of soluble and membrane-anchored OpuAC

In order to quantify the ligand affinity of free OpuAC, in comparison to the physiological situation, where the protein is covalently tethered to the OpuA translocator (TMDs), we performed a glycine betaine titration (Figure 3). For this, we used OpuAC (367C-423C) with a slightly altered labelling scheme as compared to the data presented in Figure 2, i.e., OpuAC (360C-423C). The results were qualitatively similar and only the absolute FRET efficiencies were distinct between the experiments due to differences in the inter-cysteine distances (compare apo/holo in Figure 2a and Figure 3a). By fitting the FRET efficiency histograms with two Gaussian distributions, the relative populations of the open and closed states were obtained as function of glycine betaine concentration [GB] (Figure 3b). The data points were fit to the function rc = [GB]/(K_d_ + [GB]), which results in a dissociation constant K_d_ of ~6 μM for free OpuAC (95% confidence interval); the fit is displayed as a black line in Figure 3b. In the equation rc is the fraction of OpuAC in the closed state.

**Figure 3.**
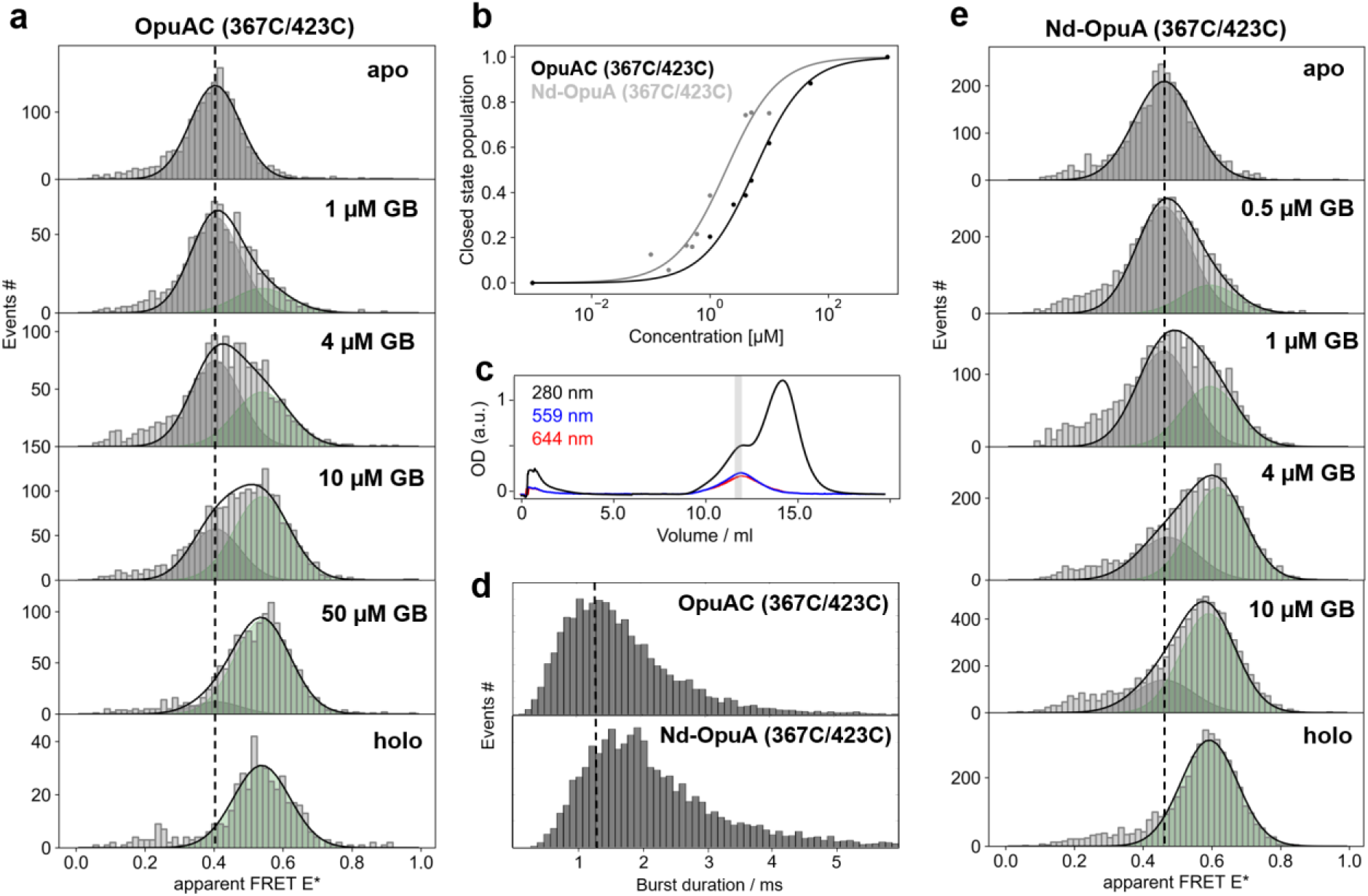
smFRET studies of intramolecular SBD conformation of free OpuAC (367C/423C) (a/b) or in the context of the transporter embedded in nanodiscs (c-e): Nd-OpuA (367C/423C). **(a/e)**. Apparent FRET efficiency histogram of OpuAC/Nd-OpuA at different glycine betaine concentrations [GB] as indicated in the Figure. The relative populations of open/apo and holo/closed state were determined using a double-gaussian fitting model with a fixed mean and width and were plotted in **(b)** as a function of ligand concentration. **(c)** Size exclusion chromatogram of fluorophore labelled Nd-OpuA (367C/423C), light grey bar annotates the fraction used for the smFRET experiments. Examples of the full data sets are shown in Figure S2/S3. **(d)** Differences in burst duration, related to size increase in Nd-OpuA, were determined from the example data sets of panel **(a/e)**.

To study the conformational switching for OpuAC within the whole transporter, we reconstituted the corresponding cysteine variant of OpuA (367C/423C) into nanodiscs using previously established protocols (details see Material and Methods)[40]. The experimental procedure is schematically sketched in Figure 4. In brief, OpuA derivatives were expressed and purified as described previously [40](Figure S4). The cell pellets were disrupted, and cell lysates fractionated by centrifugation to obtain membrane vesicles containing the overexpressed *L. lactis* OpuA derivatives (Figure 4, V). OpuA was solubilized with DDM and immobilized on a Ni^2+^-sepharose™ resin (Figure 4, I). Fluorophore labelling was performed at this stage. After removal of excess dye, OpuA was reconstituted into nanodiscs. The nanodiscs were purified by Size-Exclusion Chromatography (SEC) (Figure S4). Selected fractions from SEC were analysed by SDS-PAGE to verify that the transporter complex was reconstituted intact (Figure S4). Proper biochemical activity was verified by assessing the glycine betaine and potassium-induced ATPase activity, which was stimulated 4-10 fold over the basal activity (Figure S5, wt OpuA).

**Figure 4.**
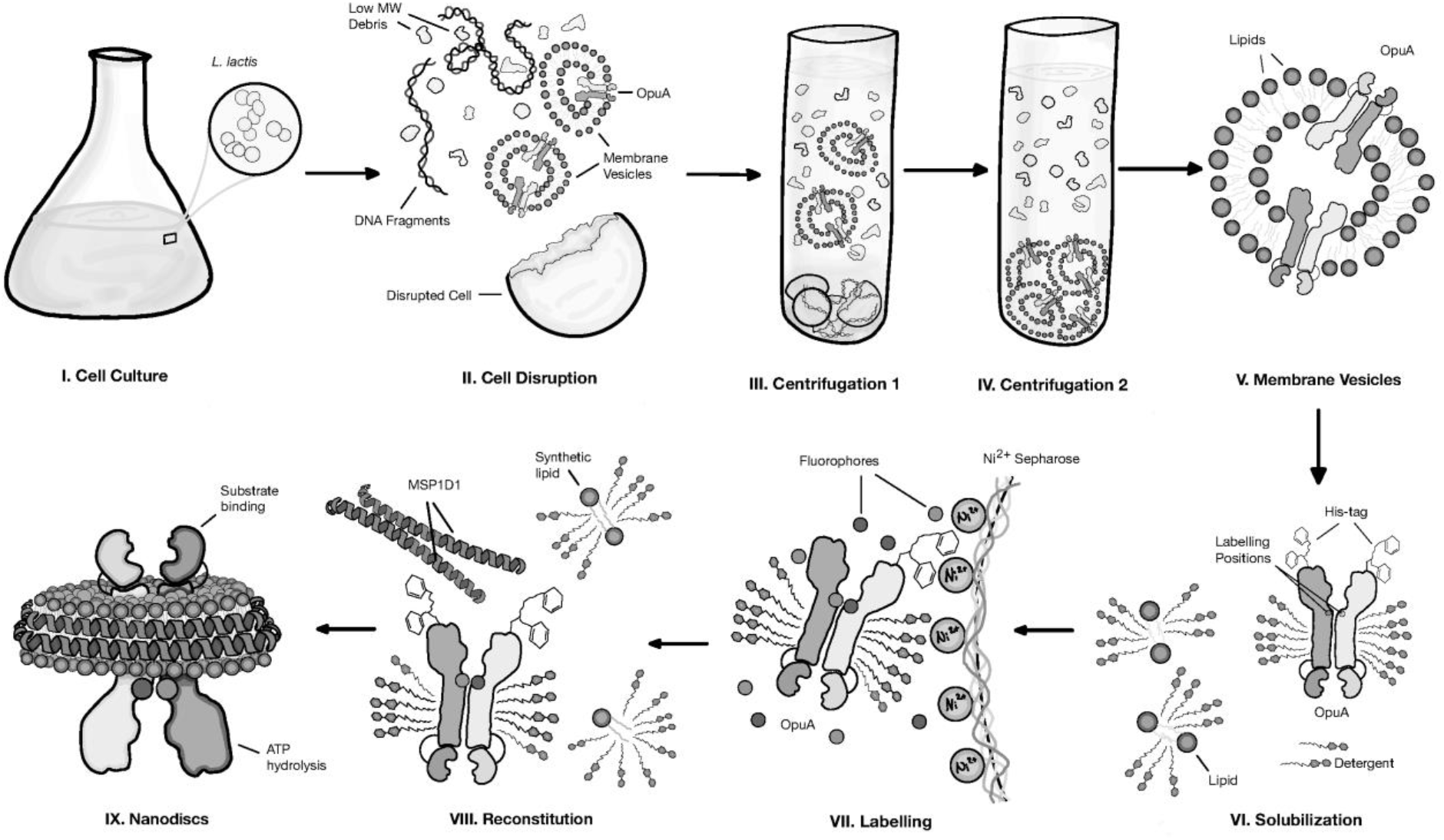
Workflow for labelling of nanodisc-reconstituted OpuA for smFRET studies. The OpuA cysteine derivatives (example used here refer to TMD positions) were overexpressed in *L. Lactis* (step I). The cells were mechanically ruptured (step II) and transporter-containing membrane vesicles were isolated (steps III-IV). At this stage the membrane vesicles can be stored at −80·C until needed for a period of 6-12 months. OpuA was released from membrane vesicles with DDM (step VI). The solubilized transporter complex was then purified and labelled utilizing affinity chromatography (step VII). In the final step, OpuA was reconstituted into nanodiscs using synthetic lipids and an amphipathic scaffold protein MSP_1_D_1_ [47] (step VIII).

On SEC, we obtained a peak eluting at ~12 ml containing the reconstituted Nd-OpuA (367C/423C) derivative [40] and a second one with the empty nanodiscs at ~14 ml (Figure 3c). By determining the absorbance at the indicated wavelengths in the corresponding elution volumes (grey bar, Figure 3c), we estimated the labelling efficiencies, i.e., the relative concentration of donor- and acceptor dye to OpuA (for details see methods parts below). As expected, from the size difference between free and nanodisc embedded OpuAC within the entire transporter, the nanodisc-reconstituted OpuA shows a slightly shifted burst-length distribution with respect to free OpuAC, indicating slower diffusion (Figure 3d).

Next, we performed smFRET on nanodisc-reconstituted Nd-OpuA (367C/423C) (Figure 3e) at different ligand concentrations, as previously done for free OpuAC (Figure 3a). OpuA can be labelled with more than 2 fluorophores in the 367C/423C variant, since four cysteines are present due to the fact that each protomer contains an OpuAC SBD. To exclude interdomain artefacts in our smFRET experiments, we analysed only molecules with a stoichiometry value >0.5 to bias our analysis towards OpuA-molecules bearing fewer fluorophores. This selection was based on the analysis of photon-counting histograms (Figure S3) of Nd-OpuA (367C/423C). We noted that the acceptor-based acceptor-emission AA was lower in the high stoichiometry region, which suggests fewer acceptor labels here. Furthermore, low S molecules also show stronger donor-quenching, which is likely due to the presence of multiple acceptor fluorophores.

In line with this interpretation and data selection, the resulting FRET efficiency histograms of soluble OpuAC and Nd-OpuA (367C/423C) were similar. In addition, the fit of the glycine betaineresponse suggests a K_d_ value of ~2 μM for Nd-OpuA (367C/423C) (Figure 3b, grey fit line), which is 3-fold higher as compared to the soluble OpuAC (~6 μM). This difference in K_d_ can be seen by distinct ratios of open/closed-state population at similar concentrations of glycine betaine (Figure 3a vs. Figure 3e), an observation that is in line with a proposed K_d_-increase for OpuAC in the context of the transporter from ref. [46]. We can thus conclude that the conformational changes of OpuAC and ligand binding are slightly influenced by its linkage with the translocator domain of OpuA.

In type-I ABC importers, TMDs are known to transit from the inward-facing state (“resting”, free or ADP-bound), to the outward-facing ATP-bound state during transport [48]. To understand how these conformational changes are transmitted to the SBDs, we monitored the conformational states of OpuAC within the nanodisc embedded OpuA (Figure 5a, Figure S6).

**Figure 5.**
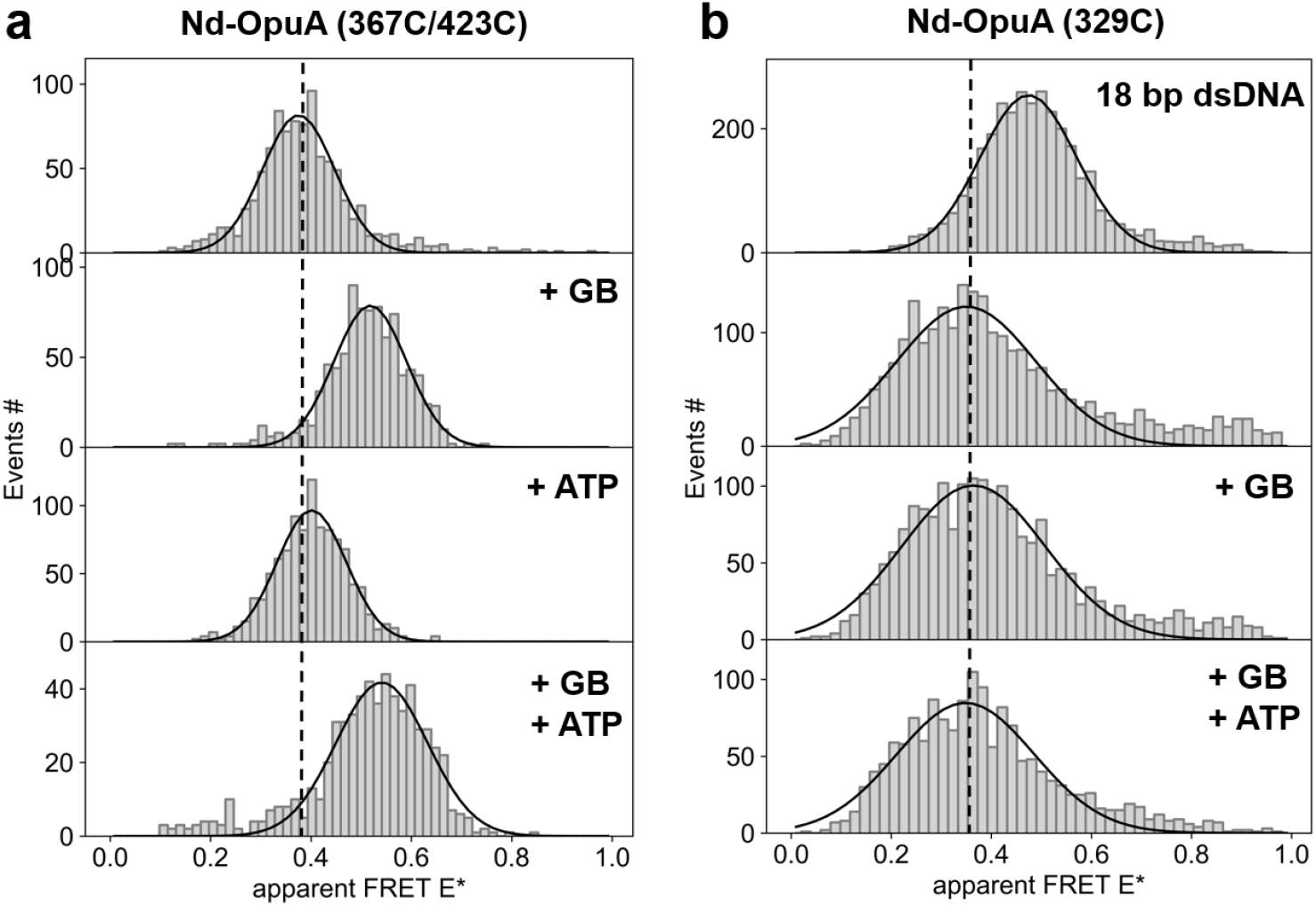
ATP- and ligand-dependent conformational switching of OpuA in nanodiscs with intramolecular SBD labels. (**a**): **Nd-OpuA (367C-423C) and interdomain labels (b): Nd-OpuA (329C)**.

Our data suggest that addition of ATP does not influence the conformational state of OpuAC either in the absence or presence of glycine betaine, using variant Nd-OpuA (367C/423C) (Figure 5a). Note that we assume here that ATP-hydrolysis is slow at room-temperature and can be used to induce conformational switching in ABC transporters using ALEX microscopy as shown previously for McjD[23]. This idea was supported by the fact that we obtained identical results as shown in Figure 5 (+ATP) with AMPPNP (Figure S6b). We thus conclude that the conformational changes of the TMDs driven by ATP binding are not transmitted to the SBDs in a way detectable by our FRET assay. However, we must note that the SBD-labelling scheme we used might impair the docking of OpuAC to the TMDs. Thus, future studies should focus on heterodimer expression of OpuA to avoid the presence of more than one donor-acceptor pair and consider backside labelling of OpuAC to avoid such complications (see discussion for details).

### The conformational arrangement of membrane-anchored SBDs

We next used the established fluorophore labelling procedure of nanodisc-reconstituted OpuA to study relative changes in interdomain distances between the two SBD domains within the full transporter (Figure 1c). For this, the labelling scheme was tailored to maintain only one cysteine per SBD and thus two per full transporter, i.e., Nd-OpuA (329C). The FRET efficiency histograms of Nd-OpuA (329C) were obtained in the apo state or in the presence of glycine betaine and/or ATP (Figure 5b). The FRET-efficiency distributions were wider as compared to a static DNA sample with one defined distance or conformational state (Figure 5b). Similar observations were also made with another labelling positions, i.e., Nd-OpuA (440C) and Nd-OpuA (458C) displayed in Figure S6c. In comparison to the dsDNA sample, this suggests that either fast distance fluctuations occur on the sub-millisecond timescale or reveal the existence of multiple distances that give rise to overlapping distributions. To rationalize these interpretations, i.e., the existence of more than one possible structural state or a larger ensemble of states, we analysed a static DNA sample (Figure 5b, upper panel). Here, the donor-acceptor distance is 18 bp suggesting 6.1 nm distance between donor-acceptor attachment points using a linear DNA-model and 7.2 nm for cylindrical DNA model[49]. As was shown previously, the width of such a distribution depends solely on photon statistics (and background) and are characteristic of a static conformational state[31, 50]. The broad and non-specific FRET-efficiency distributions of the interdomain labels in Nd-OpuA (329C) can also explain the lower data quality of Nd-OpuA (367C/423C) in comparison to soluble OpuAC (367C/423C). Nd-OpuA (367C/423C) showed a significantly elevated number of unspecific FRET events, which were likely related to interdomain FRET events and were found around the major population at E* ~ 0.4 of intramolecular FRET within OpuAC (see Figure S2, line 1 vs. line 2) due to the possibility of labelling both SBDs.

### Towards smFRET studies of the TMDs

As high-resolution structures of the translocator domain (TMD) of OpuA were not available to us, we performed homology modelling by using the Swiss-Model server [51, 52] (Supplementary Note and Figure S7/8) to identify suitable positions in the TMD for smFRET studies. The best templates for modelling were the structures of the bacterial importers for maltose (4jbw), molybdate tungstate (2onk), methionine (3dhw) and molybdate sulfate (3d31). Unfortunately, the sequence identity was <26% and the sequence similarity <32% suggesting that the resulting model may not be entirely accurate.

Keeping the limitations of the model and its unclear accuracy in mind, we focussed here on establishing an experimental pipeline to identify suitable TMD residues for smFRET with the goal to visualize changes between the inward- and outward facing conformation. As a first step, we identified non-conserved residues that were surface exposed and then residues that undergo significant distance changes between the inward and outward facing conformation in the model (Figure S7/8). Because our choices were based on a homology model, we decided to generate a larger number of cysteine derivatives (Figure 6/7a). The translocator domain of OpuA is homodimeric, and so only a single cysteine substitution per protomer would be sufficient to obtain two anchor points for fluorophores. The resulting derivatives were subsequently tested for their *in vivo/in vitro* functionality and labelling efficiency (Figure 7a).

**Figure 6.**
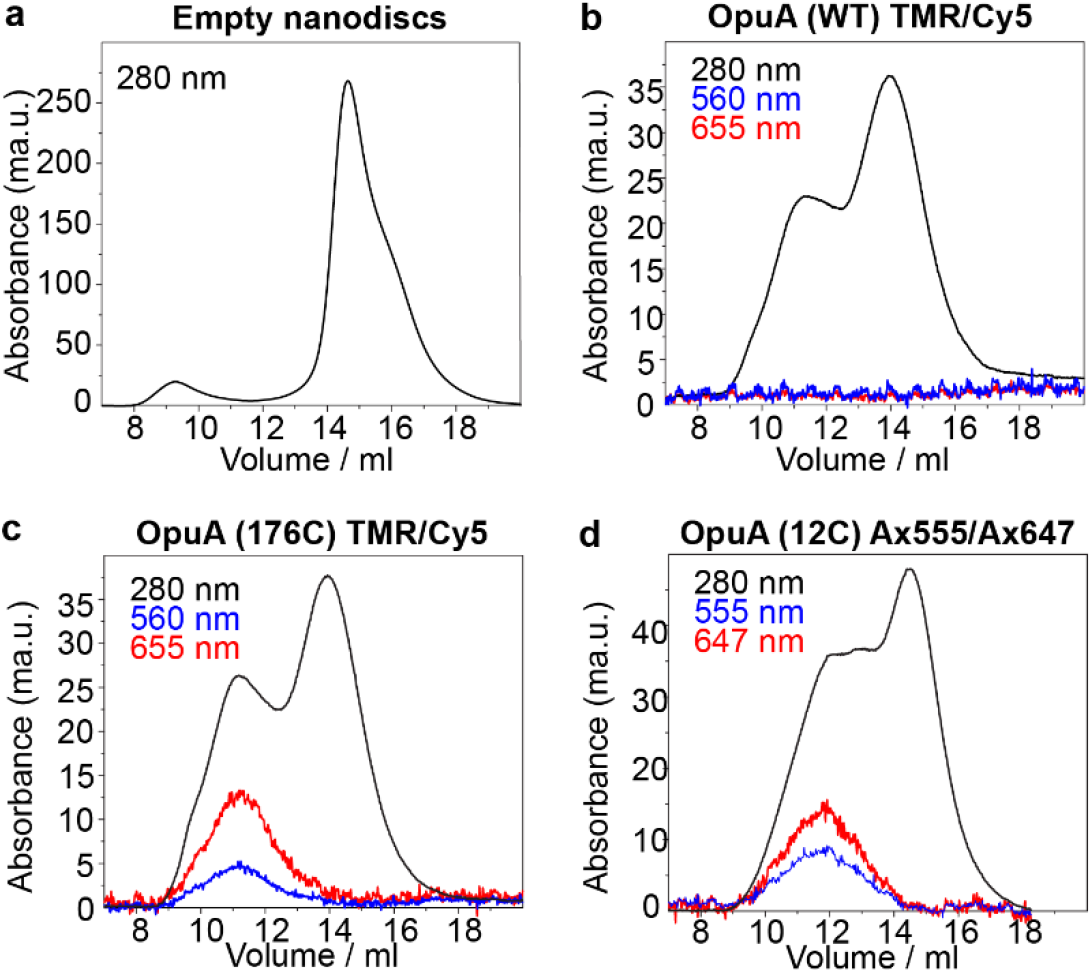
Optimization of fluorophore labelling for smFRET studies of OpuA TMD mutants. **(a)** Size exclusion chromatograms of (I) reconstituted nanodiscs without any OpuA. (b) Wildtype OpuA reconstituted in nanodiscs and labelled with TMR and Cy5 to evaluate the possibility of nonspecific binding of the fluorophores to the cysteine-less Nd-OpuA. (c): Nd-OpuA 176C labelled with TMR and Cy5. (d) Nd-OpuA 12C labelled with Alexa 555 and Alexa 647. Al nanodisc preparations show a shoulder at lower elution volumes, which might indicate nanodiscs with more than one transporter.

The selected TMD-derivatives were labelled with different fluorophore pairs as exemplified in Figure 6 for Nd-OpuA (176C) with TMR/Cy5 and for Nd-OpuA (12C) with Alexa555/Alexa647. As shown in Figure 6, wildtype OpuA was not labelled with maleimide fluorophores due to the absence of thiol-groups. The labelling efficiencies were derived by monitoring the absorbance of the protein and the fluorescent dyes at their corresponding maximum wavelength. Concentration determination of protein and fluorophores was performed using the Lambert-Beer law and published extinction coefficients (see methods section and Table S1). Unfortunately, the contribution of MSP_1_D_1_ to the 280 nm absorbance could not be deconvoluted, because methods to perform such a deconvolution only became available quite recently [53]. The labelled fraction (~11-12 ml) expected to contain the two OpuA components in equimolar ratio, was selected for subsequent testing in biochemical and biophysical assays (Figure 6c, d).

To test the effect of the cysteine mutations on function, we first performed *in vivo* uptake assays using radiolabelled glycine betaine (Figure 7a) as described in ref. [40]. Moreover, to evaluate the effect of fluorophore attachment, we determined ATP hydrolysis rates for OpuA (12C) labelled with TMR-Cy5 pair in comparison to OpuA wildtype. In Figure 7a, we summarize the relevant, substrate-stimulated ATPase activities relative to wildtype; full data sets of different biochemical conditions (apo, KCl, GB and KCl) are shown in Supplementary Figure S6. We identified the OpuA (12C) derivative as a promising candidate with a good labelling efficiency (~40% total labelling efficiency of donor and acceptor dye) and ~70% substrate-stimulated ATPase activity when labelled.

We performed ALEX experiments on the apo-protein state of OpuA (12C) under ATP- and ligand-free conditions in both detergent and nanodiscs (Figure 7c). Discrepancies were observed between labelling efficiency of OpuA in detergent and labelling efficiency of OpuA in nanodiscs (Figure 7c) due to the biobeads interacting with the fluorophores (Figure S9, Methods: Fluorophore labelling of OpuA). For nanodisc-reconstituted OpuA the labelling efficiency was ~40% for TMR-Cy5 (Figure 6c), the donor-acceptor fraction was expected to be <10%. It is likely for this reason that in our preliminary ALEX experiments we observed a high degree of random coincidence, i.e., coincident detection of donor- and acceptor-only species due to the high concentrations needed to see FRET events. This random coincidence was observed as a smear from low E/high S to low S values in this data set (Figure 7c, l-shaped lines between donor- and acceptor-only at low FRET efficiency). Nevertheless, for TMR-Cy5 labelled protein, there was a clear species at E* ~ 0.7. The presence of low-FRET species representing a more open conformational state of the protein hidden by the high level of random coincidence cannot be ruled out. All other cysteine variants (Figure 7a) gave smFRET results with still smaller donor-acceptor yields and thus no interpretable FRET-efficiency histograms.

**Figure 7.**
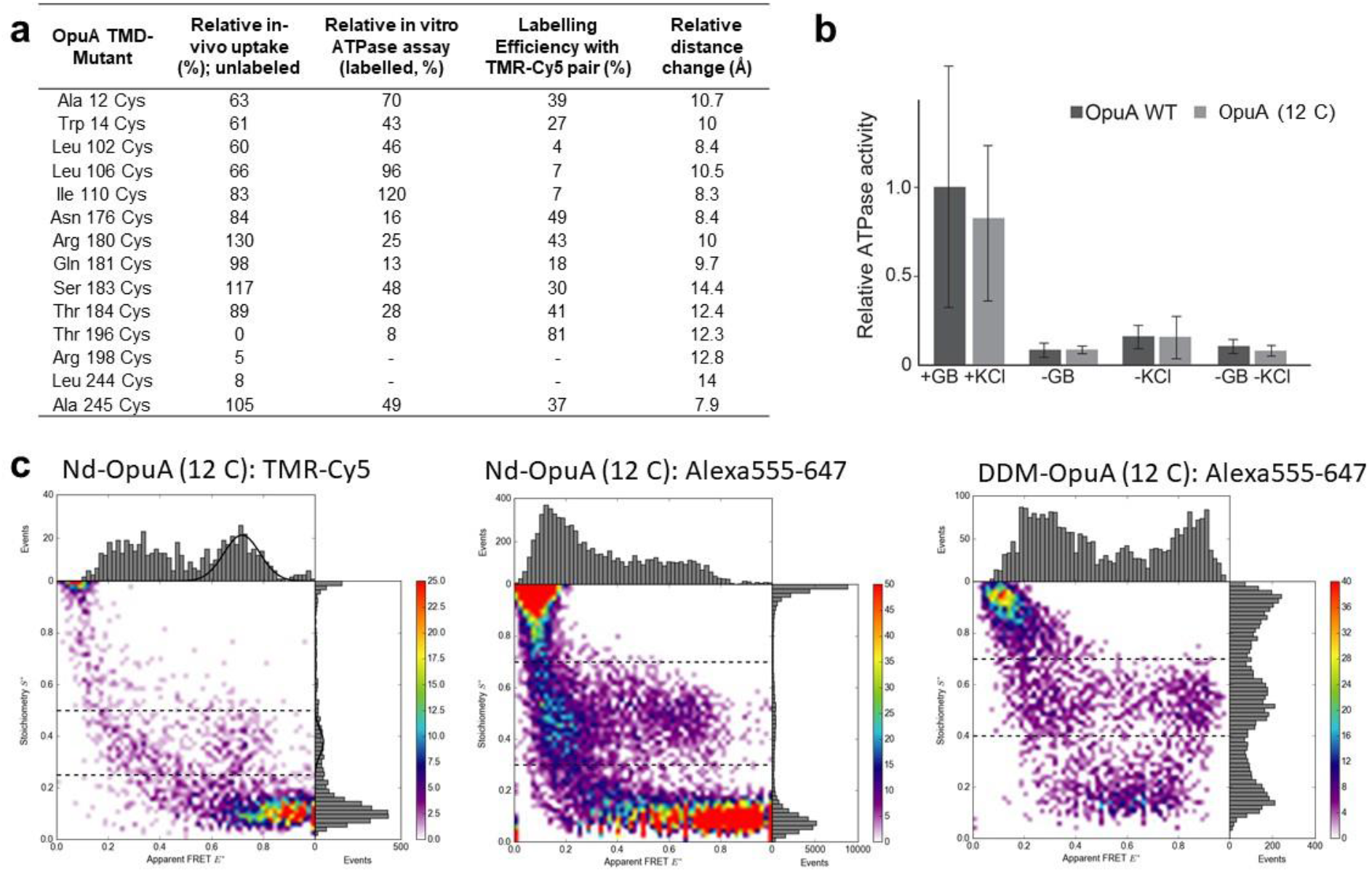
smFRET studies of TMD-labelled OpuA with OpuA (12C). **(a)** To select the appropriate cysteine derivatives for smFRET experiments we tested the effect of the cysteine point mutations and the fluorophore labelling on the activity of OpuA. Labelling efficiency was calculated by considering both donor and acceptor-labelling yields from SEC runs when comparing protein and dye absorbance. In the last column we also present the distance change between the modelled inward and outward conformation, an important parameter for the sensitivity of the smFRET assay. **(b)** ATPase activity comparison between labelled Nd-OpuA (12C) and OpuA wiletype (WT); mean ± SEM is shown (N = 3). **(c)** Nd-OpuA (12C) with different fluorophore pairs showing increased donor-acceptor yield for use of Alexa555-647. For the data all photon burst-search was used (M = 15, T = 500 μs and L = 50) with additional thresholding of all photons >100. The S-range for 1D-E* histograms is indicated in the figure as dashed line.

We next tried to use a commonly employed fluorophore pair Alexa Fluor 555-647 and obtained labelling efficiencies of 39% in nanodiscs and 46% in detergent (Figure 7c). Despite the fact that the fluorophores perform better in terms of brightness, Nd-OpuA (12C) labelling with Alexa dyes resulted in noisy data in both nanodiscs and detergent. Based on the quality of the available data no useful interpretation was possible. We attribute the poor quality of the resulting histograms to artefactual interactions of the dyes with the protein, e.g., the aggregation of OpuA in detergent might have occurred and could explain why higher yields of donor-acceptor containing molecules found in the detergent solution in comparison to lipid nanodiscs (Figure 7c). This example shows clearly that better data quality is required to rule out artefacts and the availability of a high-resolution structures would facilitate assay design and final interpretation of smFRET results in a meaningful way.

## DISCUSSION AND CONCLUSION

Long-distance allosteric communication is central to the function of ABC importers[54] and exporters[55]. The association and dissociation events of the NBDs that are driven by ATP binding and hydrolysis are transmitted to TMDs, which undergo conformational rearrangements that facilitate transport. SBDs act as primary receptors to bind and donate their respective substrates for transport initiation. The tight connectivity between the different domains is undisputed and its functional repercussions have been verified in many systems, e.g., the means by which the conformational changes of SBDs dictate substrate specificity[21] and transport[19]. However, many salient features of these interdomain associations remain elusive and the latter form the basis of the distinction between different transporter systems.

Substrate capture by SBDs is the initial step for transport. In type-I systems, the SBDs alternate between two distinct conformations, an open unliganded form in which the two lobes resemble an open book, and a closed liganded one, in which the ligand is captured by the well-known Venus fly trap motion[56]. In contrast, in type-II importers the structural rearrangements in the SBDs driven by the ligand are minor[56]. OpuAC from *L. lactis* is a type-I SBD linked to the TMDs. Thus, two OpuAC domains are present per functional transporter. In this paper, the SBD OpuAC was labelled with donor and acceptor fluorophores and its conformational states were probed by smFRET. Soluble OpuAC, as also previously reported for other members of the periplasmic binding protein family[21], binds its substrate by the Venus fly trap motion and most likely an induced fit mechanism according to our data. Its structural transitions are ligand dependent, and by titrating the ligand we derived K_d_ values by kinetic rate analysis and occupation ratios of the conformational states. These values are in excellent agreement with the ones determined in bulk, indicating that labeling and our smFRET assays do not interfere with ligand binding.

We also probed the conformational changes of OpuAC within the full transporter complex. For this, we tested many distinct labelling protocols including fluorophore addition (i) during the purification, (ii) solubilization or (iii) reconstitution of the entire transporter. The highest labeling efficiency could be achieved when the fluorescent dyes were added to detergent-solubilized OpuA that was immobilized on the Ni-NTA column material (Figure 4). The 2D ALEX plot of the labeled OpuAC within OpuA indicated the existence of multiple labelling populations, as expected because of the two OpuAC domains per transporter, and the possibility of attachment of two fluorescent labels per OpuAC. With analysis based on fluorophore brightness (Figure S3), we can separate the populations (Figure 1a/3). The results indicate that OpuAC within the entire transporter samples similar conformation states: open-unliganded and closed-liganded state. Furthermore, ligand titrations indicated that OpuAC within OpuA might have elevated binding-affinity for glycine betaine. We must note that currently we cannot exclude that the labels hinder the docking process and further optimization of the labelling scheme is required to minimize this possibility.

The second step of transport is represented by the interaction of the SBD with the TMDs. In type-I importers, the TMDs differentiate between the open-unliganded or the closed-liganded state of the SBD. It is believed that interaction of the closed liganded state with the TMD [19] efficiently triggers ATP hydrolysis[42, 57]. Conversely, the ATPase activity of type-II importers can be ligand independent or shows at least much smaller stimulation values in the presence of liganded SBD/SBP[58, 59]. Such findings suggest a communication of the SBDs and the NBDs via the TMDs. Within our assays, we probed the OpuAC conformational states during transport conditions (glycine betaine + ATP) and throughout its resting state (apo). Addition of ATP and glycine betaine had no effect on the conformational dynamics of OpuAC. Since a single SBD interacts with the TMD to deliver one substrate per transport cycle [42], an important question is whether the two OpuAC domains interact stochastically or in a concerted fashion with the TMD to release the substrate. To probe the relative movement of the two OpuAC domains, we placed one probe per domain using three different OpuA derivatives (329C, 440C, 458C). Our results indicate that probably, there are fast, non-concerted motions between the two SBDs that are independent of the state of the TMDs, since we did not find changes in the distribution upon addition of ATP/GB. All three mutants also gave fairly broad featureless distributions (Figure 5b/S6c) which might relate to uncorrelated stochastic motion. Future studies should reveal the relative motions of the two OpuAC domains during nucleotide cycling in more detail using advanced fluorescence methods with improved temporal resolution such as multiparameter fluorescence detection (MFD analysis[30]), fluorescence correlation spectroscopy or pulsed-interleave excitation[50].

The absence of ATP/ligand-driven effects in our study might, however, also result from a failure of our labelled mutants to dock to the translocator TMDs due to steric hindrance effects and the chosen positions, which were only optimized for smFRET and not the biochemical activity. In future experiments, such effects must be avoided by use of alternative labelling schemes, e.g., on the back of OpuAC. Optimal labelling residues on the back of OpuAC (hinge region) for use of Cy3B/ATTO647N or Alexa555/647 would be A414C-A569C, A414C-A565C, A414C-A570C or A414C-A562C for intradomain monitoring (or any of these single cysteines for interdomain monitoring) considering the Förster radii of the two fluorophore pairs and the fact that they were used successfully for labelling nanodisc-reconstituted OpuA here.

The next step of transport involves the TMD conformational changes that needs to occur to promote the passage of the substrate to the cell interior. Those changes are assumed to be strongly coupled to the ATPase cycle in type-I importers [54]. Type-I ABC transporters switch from the resting state of the inward facing conformation to the outward, a process coupled to ATP and SBD binding events – which is distinct for type-II importers. That might be the reason that in type-I importers the ATPase activity is better regulated (i.e., here ATP hydrolysis is generally well coupled to translocation), whereas type-II manifest high basal ATPase activities in the absence of SBD and substrate. To probe transitions between the inward and outward facing conformation of OpuA, we produced a large number of cysteine derivatives. We started with a set of 14 mutants of which 11 still proved to be functional (>60% transport activity *in vivo).* Remarkably, while the chosen positions are surface-exposed according to our homology model, the majority could not be labeled well with dyes, suggesting that the position might be distinct from what is suggested in the model. The low sequence conservation of OpuA with available structures renders this finding not surprising. However, we selected the best derivative with respect to the labeling efficiency and the retention of the *in vivo* and *in vitro* functionality. The interpretations of the smFRET results of Nd-OpuA (12C) are still not fully clear since the resulting donor-acceptor yields were too low for in depth interpretation. Without further studies we cannot claim with confidence whether we observe one or multiple conformational states at this position and also whether the choice of different fluorophore-pairs might have impacted our observations (Figure 7c). In the future, more experiments with improvements on labelling efficiency and in particular donor-acceptor-containing functional transporter molecules are required. This type of problem, i.e., high labelling efficiency yet low donor-acceptor fractions when labelling TMD-residues in ABC transporters, was reported by us [23] and others [24] before. This complication was much less pronounced when the labelling was performed on the NBDs [23]. Improvement of this aspect could reduce the measurement times from >1 hour for proper statistics, to much shorter periods and would allow further testing of different biochemical studies of the transport cycles.

In summary, despite all problems encountered, we here pave the way for future experimental smFRET studies that aim at understanding the salient features of inter- and intra-domain communication in ABC transporters and in particular for OpuA.

## METHODS

### OpuA mutagenesis, expression and membrane vesicle isolation

The OpuA nucleotide sequence, (no endogenous cysteines), from *Lactococcus Lactis* was subcloned to the pBR322 vector (Addgene). The plasmid was used for the introduction of point cysteine mutations eligible for labelling by QC-PCR (Supplementary Table S2). The point mutants were subsequently cloned in the pNZopuAHis plasmid (C-terminal 6-HIS-tag) using *EcoRV-AlnwnI* (NEB) for the substrate-binding domain mutants and *βam*HI-*Alwn*I (NEB) for the transmembrane domain mutants (Appendix). For the expression of the mutant proteins *L. lactis* Opu401 strain was used, which contains deletion of endogenous *OpuA* genes from the chromosome. 2-5-liter cultures were grown anaerobically at 30 °C in 2% (w/v) Gistex LS (Strik BV, Eemnes, The Netherlands) and 200 mM potassium phosphate (KPi), pH 7.4, supplemented with 1.0% (w/v) glucose and 5 μg/ml chloramphenicol. At OD_600_ ~2 the *nisA* promoter was induced by the addition of 1 ng/ml nisin. Two hours later the cells were harvested by centrifugation (6000 *g;* 15 minutes; 4°C) and stored at −20°. For the isolation of *L. lactis* membrane vesicles all handling was done at 4°C unless stated otherwise. The harvested pellet was resuspended in 50 mM KPi, 200 mM KCl, 20% glycerol, pH 7.4 (buffer A) in the presence of 1.5 mM dithiothreitol (DTT). To reduce the viscosity caused by the release of DNA, 100 μg/ml DNase and 2 mM MgSO4 were added. The suspension was disrupted through a continuous disruption cycle, twice at 40 Psi in a Constant Cell Disruption System LTD. In the disrupted material 1mM PMSF and 5mM EDTA, pH 8.0 was added. First, large cellular debris were removed by centrifugation (11,800 *g;* 20 minutes; 4°C) the pellet was discarded and the supernatant was centrifuged again (125,000 *g;* 60 minutes; 4°C). The pellet containing membrane vesicles was resuspended in buffer A and 1mM DTT and centrifuged again (125,000 *g;* 60 minutes; 4°C) to remove any remaining soluble components. The isolated membrane vesicles were resuspended in buffer A in the presence of 1mM DTT, aliquoted and flash frozen in liquid N2 and then stored in −80°C. The total protein concentration was determined using the Pierce™ BCA Protein Assay Kit.

### OpuA purification

All the handling described below was done at 4°C unless mentioned otherwise. The stored membrane vesicles were thawed and resuspended in buffer A (50 mM KPi, 200 mM KCl, 20% glycerol, pH 7.4) with the addition of 1 mM DTT and 10 mM *n*-Dodecyl β-D-maltoside (DDM) and incubated for 60 minutes while gently agitated. The solubilized protein was separated from any insolubilized material by centrifugation (267,000 *g*; 20 minutes; 4°C). The harvested supernatant was diluted with buffer A to a final DDM concentration of 2 mM and loaded to Ni^2+^-Sepharose™ 6 fast flow resin (GE Healthcare, already equilibrated with 10 CV buffer A with 780 μM DDM) and was incubated for 60 minutes under gentle agitation. The resin-bound material was washed three consecutive times (10 CV of each: buffer A with 1mM DTT and 780 μM DDM, buffer A with 1 mM DTT and 20 mM imidazole and 780 μM DDM, and buffer A with 1 mM DTT and 40 mM imidazole and 780 μM DDM) to remove all the weakly bound proteins. The purified protein was then eluted with buffer A with 200 mM imidazole and 780 μM DDM.

### OpuA nanodisc formation and purification

An optimized synthetic lipid mixture dissolved in 50 mM KPi, pH 7.0 (50% 1,2-dioleoyl-sn-glycero-3-phosphoethanolamine, DOPE: 12% 1,2-dioleoyl-sn-glycero-3-phosphocholine, DOPC: 38% 1,2-dioleoyl-sn-glycero-3-phospho-(1’-rac-glycerol), DOPG, Avanti Polar Lipids) was prepared as described [40]. The lipid mixture was thawed and subsequently extruded through a 400 nm pore size polycarbonate filter, generating large unilamellar vesicles. 12 mM DDM was added and the mixture was vortexed until optically clear. The final reconstitution reaction was 50 mM KPi, 20% glycerol, 12 mM DDM and contained 1 nmol OpuA (labelled/unlabeled), 10 nmol MSP_1_D_1_, 1 μmol lipids in a final volume of 700 μl. To control for size and formation of nanodiscs, reconstitution reactions without OpuA were carried out and tested using size-exclusion chromatography (SEC). The reaction was incubated for 60 minutes at 4°C, while gently agitated. Then, 500 mg of SM2 bio-beads was added to the reaction volume and incubated for 1-12 hours with best result at 1-2 hours of incubation. The bio beads where removed and the solution was centrifuged (18000 *g;* 10 minutes; 4 °C) to precipitate any aggregated lipids and proteins. To determine the composition of the nanodiscs and to separate the nanodiscs from aggregates and empty nanodiscs, the supernatant from the previous step was purified using SEC, using a Superdex 200 10/300 GL column (GE Healthcare) that was previously equilibrated with 50 mM KPi, pH7.0, 200 mM KCl, 4% w/v glycerol buffer. The protein composition of the SEC fractions was verified by SDS-PAGE on 12% polyacrylamide gels.

### MSP_1_D_1_ expression and purification

Membrane scaffold protein MSP_1_D_1_ was used for the formation of nanodiscs [40]. *E. coli* BL21(DE3) cells were freshly transformed with the pMSP_1_D_1_ plasmid and grown in 2 l Terrific Broth-kanamycin (10 μg/ml) medium at 37 °C under aerobic conditions. At OD_600_ of 1.5 the culture was induced with 1 mM isopropyl 1-thio-D-galactopyranoside (IPTG), three hours later the cells were harvested by centrifugation (8000 g; 20min; 4 °C). From this step onwards all the processes were done at 4°C and all solutions were at 4°C, unless stated otherwise. The cell pellet was re-suspended in 20 mM KPi, 1% Triton X-100, 1 mM PMSF at pH 7.4. The cells were lysed by sonification (5 sec on/ 5 sec off, 70% amplitude, 3 minutes). The cell lysate was fractionated by centrifugation (125000 g; 75 min; 4°C) and the pellet was discarded. Ni^2+^-Sepharose™ 6 fast flow resin (GE Healthcare) was equilibrated with 10 column volumes of 40 mM KPi, pH 7.4 and the supernatant of the previous centrifugation was gravity loaded to the column. The resin-bound MSP_1_D_1_ was sequentially washed with 10 column volumes of 40 mM Tris/HCl, 0.3 M NaCl, 1% Triton X-100, pH 8.0, then 40 mM Tris/HCl, 0.3 M NaCl, 50 mM sodium cholate, 20 mM imidazole, pH 8.0 and lastly with 40 mM Tris/HCl, 0.3 M NaCl, 50 mM imidazole, pH 8.0. MSP_1_D_1_ was then eluted with 40 mM Tris/HCl, 0.3 M NaCl, 0.4 M Imidazole, pH 8.0. The eluent was dialyzed overnight (SnakeSkin™ Dialysis Tubing, 7K MWCO ThermoFisher Scientific) against 20 mM Tris/HCl, 0.1 M NaCl, 0.5 mM EDTA, pH 7.4. Lastly, it was dialyzed once more overnight against 20 mM Tris/HCl, 0.1 M NaCl, 0.5 mM EDTA, 50% glycerol, pH 7.4. The protein concentration was calculated by absorbance at 280 nm (extinction coefficient 21,000 M^-1^ cm^-1^; see Supplementary Figure S5a). The purified protein was aliquoted and stored at −20 °C.

### ATPase activity assay

To determine the activity of the reconstituted OpuA, a coupled-enzyme activity assay was used to measure the ATPase activity of the reconstituted transporter. The coupled enzymatic reaction contained ~2 units of pyruvate kinase/lactic dehydrogenase (rabbit muscle isolate mixture, Sigma Aldrich) 50 mM KPi, pH 7.0, 300 mM NADH, 4 mM phosphoenolpyruvate (PEP), 62 μM Glycine Betaine, 300 mM KCl and 4 μg of OpuA reconstituted in nanodiscs. The change in absorbance at 340 nm was observed with a Synergy MX 96-well plate reader (Bio Tek Instruments, Inc.). The reaction components were incubated in the plate reader at 30 °C for 3min. Then 10 mM MgATP, pH 7.0, was added shaking the plate for 10 seconds to mix the compounds. A seven minute kinetic read (340 nm, 30 °C) with minimal time intervals was executed. The data were corrected for the path length of each individual reaction volume. For every mutant tested, controls were done in the absence of: a) glycine betaine, b) KCl, c) glycine betaine and KCl (Supplementary Figure S6). The ATPase activity was calculated from the slope of the measurements and normalized against the WT internal control.

### Radiolabeled isotope uptake

For the in vivo radiolabeled-isotope uptake in *L. lactis* the previously published method was used [60]. Briefly, *L. lactis* Opu401 transformed with the appropriate plasmid was grown in glucose-CDM with 5 μg/mL chloramphenicol. At OD_600_ ~0.5 the culture was induced with 0.01% (v/v) nisin A (filter-sterilized culture supernatant from *L. lactis* NZ9700) until OD_600_ ~1. The cells were washed and resuspended at 2.5 mg of cell protein/ml in 50 mM HEPES-methylglucamine pH 7.3. Next, 10 mM glucose was added and the cells were incubated for 5 min at 30°C. To start the uptake a final concentration of 600 mM sucrose (osmotic stress to increase the internal ionic strength of the cell and activate OpuA), 1mM [^14^C] glycine betaine in the presence and 50 μg/ml chloramphenicol (to prevent protein synthesis) were added to a reaction volume of 500 μl. 80 μl samples were taken at regular time intervals and diluted with 2 mL of ice-cold assay buffer of equal osmolality and filtered through 0.45 μm cellulose nitrate filters under high vacuum. The membranes were washed with ice cold 50 mM HEPES-methylglucamine pH 7.3. After drying, 2 ml of scintillating liquid was used to dissolve the membranes. The radioactivity was determined in a scintillation counter.

### Fluorophore labelling of OpuA

OpuAC was labelled as described previously [21]. For the labelling process of OpuA the final elution step was eschewed during purification and the resin was drained and washed with buffer A and 780 μM DDM to remove DTT. Then 100 nmol of maleimide fluorophores (molar ratio of 7-10 fluorophores per cysteine available) were solubilized in 10 μl water-free dimethyl sulfoxide (DMSO) at room temperature and then suspended in buffer A and 780 μM DDM. The fluorophore mixture was added to the drained resin and incubated for 60min under gentle agitation. The excess of fluorophores was washed with 10 CV buffer A and 780 μM DDM and the labelled OpuA was eluted with buffer A, 200 mM imidazole, 780 μM DDM (Supplementary Figure S5b,c). The protein and fluorophore concentrations were estimated using Lambert-Beer’s law (*A = εlc).* The absorbance was calculated by measurements at 280 nm, 560 nm and 655 nm (calculating the area under the chromatogram of the relevant fraction). The path length was 0.1 cm and the extinction coefficients were available in the data sheets of the fluorophores (Supplementary Table S4). Labelling efficiency was calculated as 100 x moles of total fluorophores / moles of cysteines. The purified OpuA was directly used for reconstitution into nanodiscs.

This final protocol was obtained after we observed that labeling efficiencies dropped significantly after the reconstitution step, as we were determining the labeling efficiency of our sample prior and subsequent to the reconstitution in bilayer nanodiscs. After many trials, we could attribute this effect to be dependent on the use of SM2 bio-beads. Bio-bead systems have been used in the past to scavenge unbound rhodamine dyes [61]. To better control labelling efficiency and sticking of protein mediated via fluorophores to the bio-beads, we used MalE as test system, for which we obtained high labelling efficiency in variant T36C/S352C [21]. To see the effect of the SM2 bio-beads on a soluble protein; 2.5 nmol of MalE (T36C/S352C) was subjected to size exclusion chromatography to determine the baseline amount of protein in absorbance units (Same buffers as used for OpuA). 2.5 nmol of protein were then incubated with 1000 mg of SM2 bio-beads (the reaction volume was doubled to 1.4 ml) overnight at 4 °C (in reconstitution buffer without lipids, detergent and MSP_1_D_1_). After incubation, the SM2 bio-beads were separated from the solution by centrifugation and the supernatant was subjected to SEC. The chromatogram (Figure S9b) was corrected for the amount of supernatant lost during the separation of the SM2 bio-beads from the protein solution. Then, it was normalized against the absorbance of non-treated MalE (Figure S9a). After extended incubation times, the SM2 bio-beads can adsorb soluble proteins as indicated by a loss of ~35% of unlabeled MalE (T36C/S352C); Figure S9. To further test our hypothesis, 5 nmol of MalE (T36C/S352C) were labelled as described above with the only difference to omitting DDM. We used fluorophores Alexa Fluor 555 (Ax555, 50 nmol) and Alexa Fluor 647 (Ax647, 50 nmol) to achieve a molar ratio of 10:1 regarding flurophore and cysteine. The eluted labelled protein sample was then split in two parts. Half of it was analyzed by SEC to determine the labeling efficiency (>75%) and also provide a baseline (Figure S9c). The other half was incubated with 1000 mg SM2 bio-beads (1.4ml reaction volume) for 3 hours at 4 °C (in reconstitution buffer without lipids, detergent and MSP_1_D_1_; Figure S9c). After the incubation MalE (T36C/S352C) was separated from the SM2 bio-beads and again analyzed by size exclusion chromatography. We observed a dramatic reduction in retrievable labelled protein (Figure S9d). Based on these results further optimization was conducted, in which we reduced the duration of the nanodisc formation reaction to 1-2 h as described above with the best labelling results observed at 1h.

### OpuA structure modelling

The homology model for the prediction for candidate mutagenic sites was done using Swiss Model Server [51, 52]. The template structure used were the molybdate/tungstate ABC transporter from *Archaeoglobus fulgidus* (ModBC, 2onk), the Maltose transport system from *Escherichia coli* (MalG, 2r6g and 4jbw), the molybdate/tungstate transporter from *Methanosarcina acetivorans* (ModBC, 3d31) and the methionine importer (MetNI, 3dhw). The sequence identity between OpuA and the templates was between 20.4% and 26.6 while similarity was ~30%. To calculate the relative distant change the produced model was aligned with the inward and outward facing conformations of the maltose transporter in PyMol [62] and the distances were measured also with PyMol for Ca-positions of the residues.

### Solution smFRET measurements and data analysis

The methods and experimental devices used here have been described previously in detail [19, 20, 63, 64]. In brief, labelled OpuAC or OpuA was diluted to 20-100 pM in imaging buffer: 50mM potassium phosphate pH 7.4, 1mM Trolox and 10mM cysteamine (pH 7.5; Sigma-Aldrich). For this, the diluted protein was loaded to BSA (200 μl of 1 mg/ml for 30 seconds) treated coverslips (no. 1.5H precision cover slides, VWR Marienfeld), minimizing fluorophore interactions with the glass slide. The measurements were done using a custom-built confocal microscope[19, 20] at room temperature. Excitation was at 532 and 640 nm in accordance with the fluorophore absorbance maxima (SuperK Extreme, NKT Photonics, Denmark). Alternation between the two excitation wavelengths was achieved by 50 μs alternation. The output beam was coupled to a single-mode fiber (PM-S405-XP, Thorlabs, United Kingdom) and recollimated (MB06, Q-Optics/ Linos, Germany) before entering an oil immersion objective (60×, NA 1.35, UPLSAPO 60XO, Olympus, Germany). Excitation and emission were separated by a dichroic beam splitter (zt532/642rpc, AHF Analysentechnik, Germany) mounted in an inverse microscope body (IX71, Olympus, Germany). Fluorescence emitted by diffusing molecules in solution was collected by the same oil objective, focused onto a 50 nm pinhole and spectrally separated (640DCXR, AHF Analysentechnik, Germany) onto two APDs (τ-spad, < 50 dark counts/s, Picoquant, Germany) with the appropriate spectral filtering (donor channel: HC582/75; acceptor channel: Edge Basic 647LP; both AHF Analysentechnik, Germany). Unless mentioned otherwise in the figure legends ALEX data [23, 63] was analysed with a dual-colour burst search (M = 15, T = 500 μs and L = 25) and the resulting data were plotted with 61×61 bins considering an S-range of 0.3-0.8 with additional thresholding of all photons >100. The data of OpuA (A12C) were collected by a similar confocal microscopy setup as described in [63]. Here, the excitation was done via two Coherent Obis lasers centered at 532 and 637 nm and the objective lens was 60X, NA 1.2, UPlanSAPO 60XO (Olympus, NL).

### Surface scanning microscopy and data analysis

For surface immobilization of OpuA nanodiscs custom flow cells were made as previously described in [19, 23, 64]. The flow cell surface was functionalized at room temperature with a neutravidin solution; 0.2 mg/ml neutravidin (Invitrogen, United States) in 50 mM potassium phosphate pH 7.4, filtered with 250 μm syringe filter (buffer B) for 5-10 minutes. The unbound excess of neutravidin was washed with the same buffer. Subsequently, the surface of the flow cell was incubated with an anti-His antibody (in buffer B) for 5 minutes and washed again with buffer B. Then His-tag containing OpuAC was introduced to the flow cell (in buffer B supplemented with 10 mM of (±)6-Hydroxy-2,5,7,8-tetramethylchromane-2-carboxylic acid (Trolox; Merck) as a photostabilizer[20]) and incubated for 30-120 seconds while at the same time scanning the surface to determine the optimum density. When the particle density was adequate the excess of labelled proteins was washed away, using buffer B with 10 mM Trolox. Fluorescence traces were recorded at room temperature. The fluorescent trajectories were analysed using a hidden Markov Model[65] as described in [21]. The binning time was 1.5 ms.

## ACKNOWLEGDEMENTS

This work was financed by an NWO Veni grant (722.012.012 to G.G.), an ERC Starting Grant (ERC-STG 638536 – SM-IMPORT to T.C.) and the Zernike Institute for Advanced Materials (support to G.G. and T.C.). G.G. also acknowledges an EMBO fellowship (long-term fellowship ALF 47-2012), and the Rega Foundation postdoctoral program of KU Leuven. We thank J. Hammerl for preparation of Figure 4 and C. Gebhardt for providing data analysis scripts. We thank B. Poolman for continuous support of the project, which included conceptional discussions on the structure and function of OpuA and access to materials and protocols. We finally thank G. K. Schuurman-Wolters for experimental support, B. Poolman and D. A. Griffith for critically reading the manuscript.

## AUTHOR CONTRIBUTIONS

G.G. and T.C. conceived and designed the study and supervised the project. K.T., R.V., M.d.K and G.G., performed molecular biology and biochemistry studies. R.V. and G.G. developed the labelling protocols. K.T., R.V., M.d.B. and G.G. performed single-molecule experiments. K.T., M.d.B., G.G. and T.C. analysed data. K.T., G.G. and T.C. prepared figures and wrote the manuscript. All authors contributed to discussion of the research and approved the final version of the manuscript.

## Supporting information

**Content: Additional data and SI Note on homology modelling**

**Figure S1.**
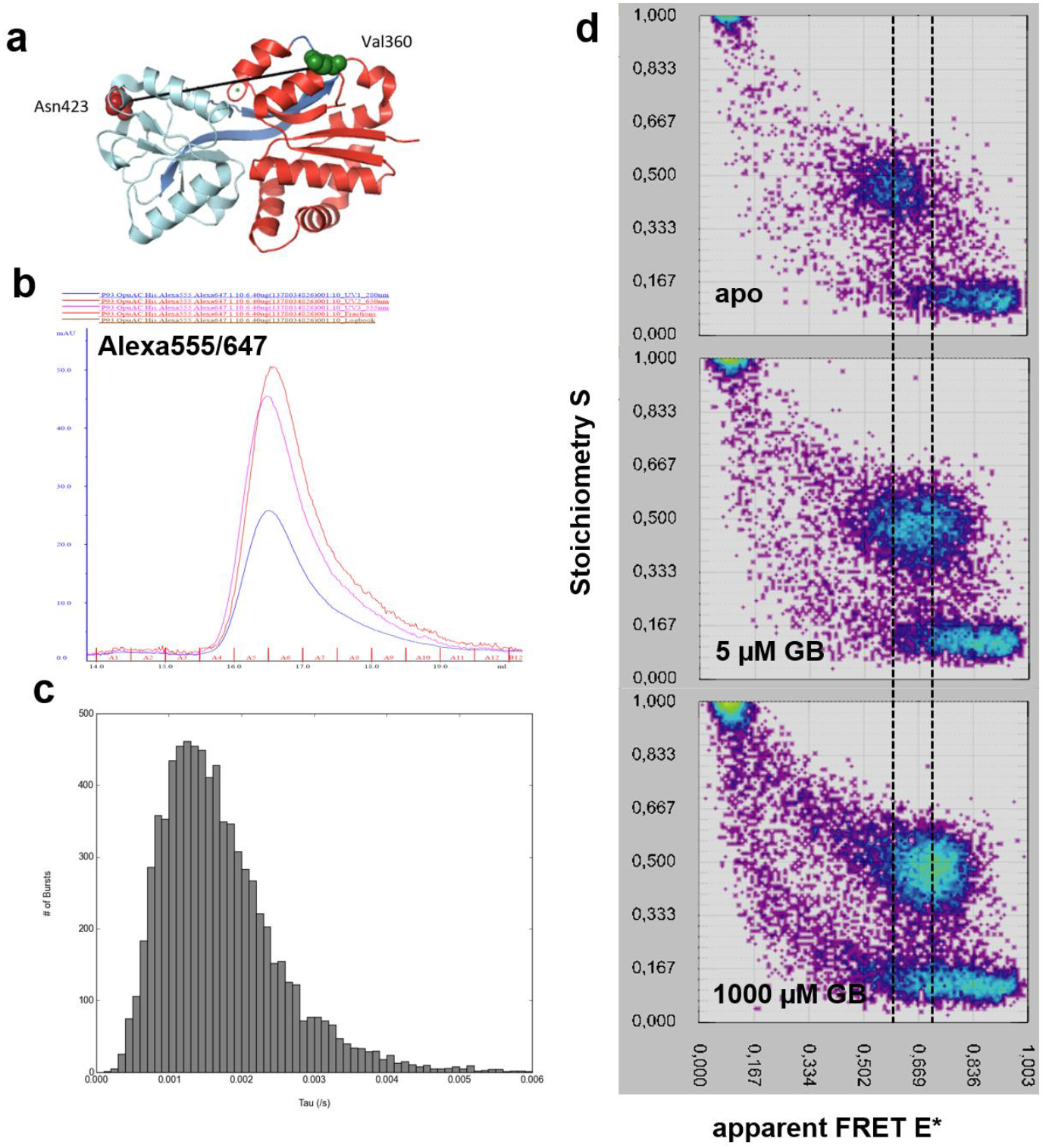
General description of smFRET experiments and analysis on the example of OpuAC (V360C/N423C). **(a)** Cartoon representation of OpuAC based on 3L6G. Residues mutated to cysteines are indicated with spheres. The two rigid domains are indicated with cyan or red color, whereas the hinge (two β strands) is shown in blue color. **(b)** After labelling with Alexa 555 and Alexa 647, OpuAC (V360C/N423C) was subjected to size exclusion chromatography, while at the same time the absorbance was monitored at three wavelengths (280 nm = blue, 555 nm = cyan, 647 nm = red) to derive labeling efficiencies. **(c/d)** Fraction A6 (eluting at ~16,7 ml, panel **(b)** was subjected to smFRET ALEX experiments, from which the burst length distribution was derived (c). The number of bursts (y axis) were plotted as a function of burst duration (x axis). **(d)** 2D ALEX histograms of single molecules (each dot is representing one labeled OpuAC (V360C/N423C) molecule which has been detected in the confocal spot) are sorted depending on their Stoichiometry. Donor-only molecules have high S > 0.8 with low E < 0.2, acceptor-only molecules of OpuAC are characterized by low S <a 0.3 and widespread E. Donor-acceptor bearing OpuAC molecules lie in between both S-regimes 0.8 > S > 0.3 with characteristic E-distributions. This S-region is the selected for data representation in the main text figures in the 1D-E*-histograms. Panel **(d)** shows the FRET-efficiency shifts of OpuAC upon addition of its natural ligand glycine betaine, GB (apo = no ligand). The population of molecules is shifting from the low FRET apo condition to the high FRET fully saturating (1000 μM of glycine betaine) conditions as indicated. For data analysis in panel **(d)** an all photon burst-search was used (M = 15, T = 500 μs and L = 50) with additional thresholding of all photons >250.

**Figure S2.**
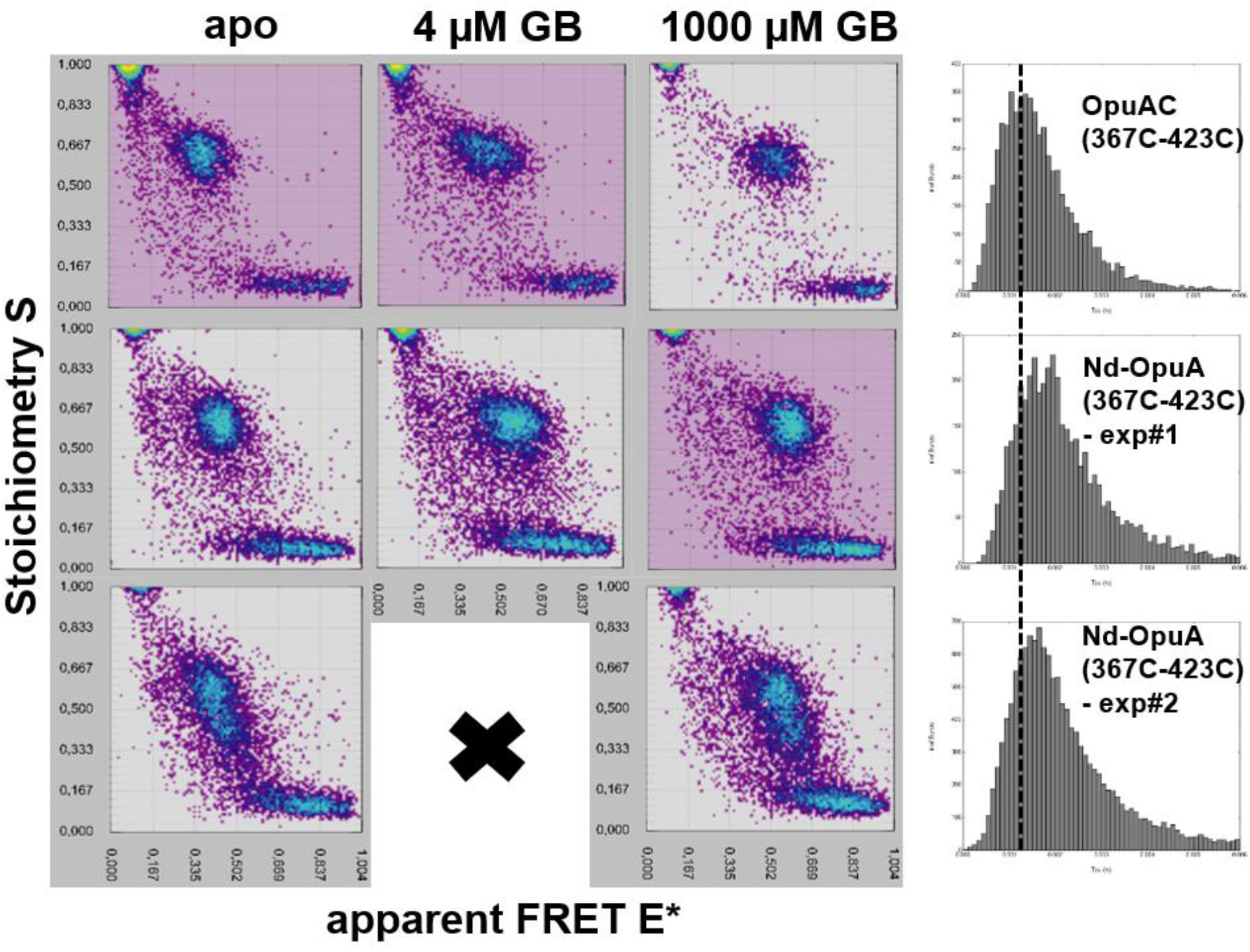
ALEX 2D plots (as in Supplementary Figure 1) of soluble OpuAC (OpuAC 367C/423C) or nanodisc reconstituted Nd-OpuA (367C/423C) labelled with Cy3B and ATTO647N fluorophores. The corresponding burst length distributions are indicated in the right part of the figure. The experiments were accomplished at apo protein, intermediate or saturating conditions of glycine betaine (GB). Free OpuAC has a narrow range of stoichiometry values 0.8 > S > 0.5, indicative of single proteins carrying one pair of single donor and a single acceptor fluorophore. A functional OpuA (Nd-OpuA (367C/423C)) consists of two OpuAC 367C/423C domains, thus four labeling positions are available per nanodisc complex. For this reason, the population in the nanodisc reconstituted OpuA has a wider S range 0.8 > S > 0.2. This is seen in different experiments with different labelling ratios of donor-acceptor and different degree of labelling (exp#1/exp2). Moreover, the burst length distribution in the nanodisc-reconstituted OpuA shifts towards higher values, indicative of a slower diffusion. In Figure S3, we tried to separate the population of molecules having only one OpuAC labelled per reconstituted transporter, from the molecules having more labels attached on both OpuAC. Importantly, the ligand binding is coupled to changes in FRET efficiency in all cases as found for undisturbed OpuAC (top row and data in the main text in Figure 2/3). For data analysis an all photon burst-search was used (M = 15, T = 500 μs and L = 50) with additional thresholding of all photons >250.

**Figure S3.**
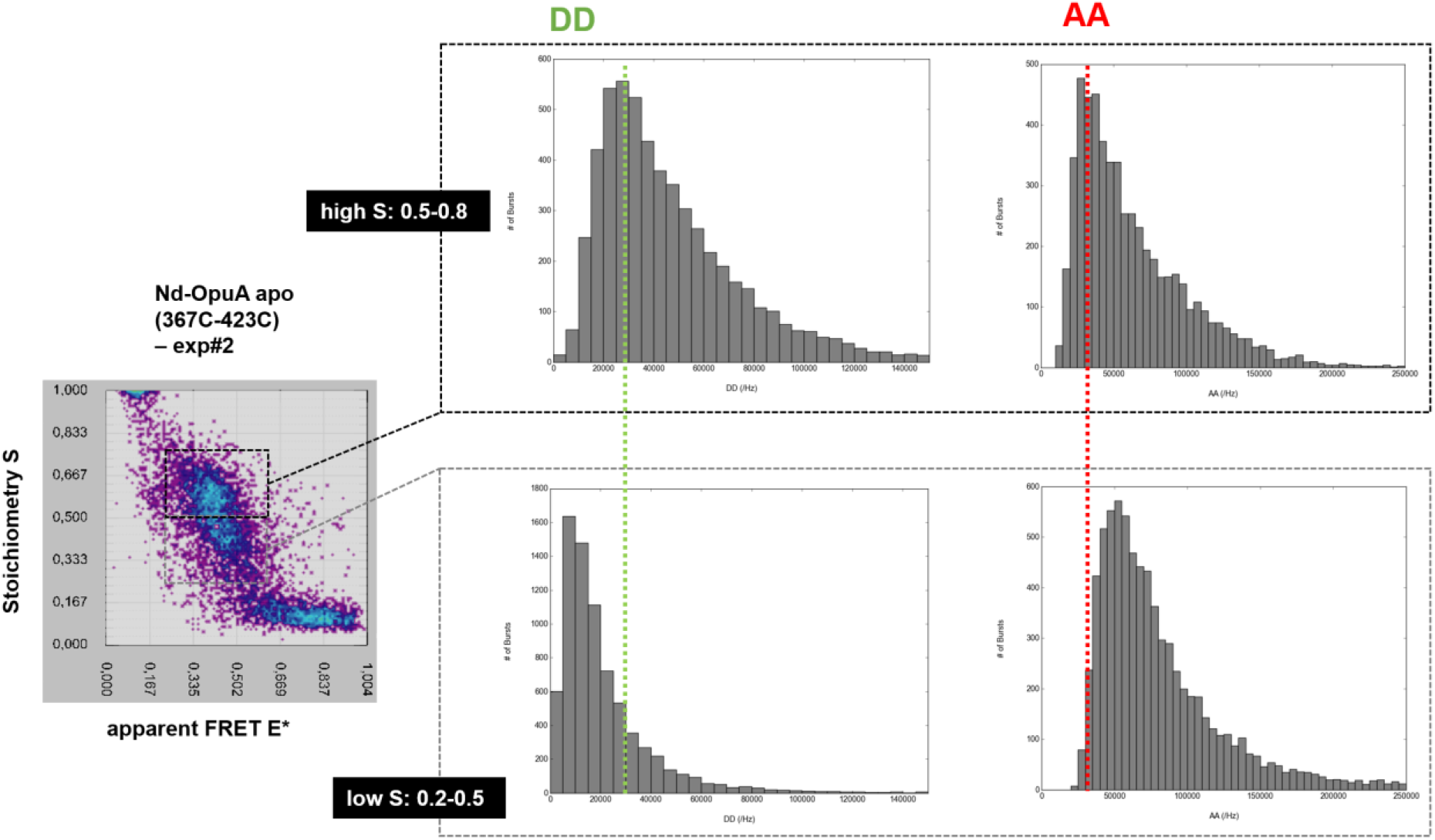
Analysis of the labelling composition of Nd-OpuA complexes. The figure shows burst-length normalized frequency of photon-count rates of donor-based donor emission (DD) and acceptor-based acceptor emission (AA). The correlation between S range (low/high as indicated) shows an increasing degree of fluorophores within the the Nd-OpuA complex. This is seen by increased mean AA-values (indicative of >1 acceptor dye) with concomitant decrease of DD-signal due to quenching via e.g., FRET. It is seen in comparison with Figure S2 that still the intermediate S-population carries useful information on the donor-acceptor distance for high S-values > 0.5 which has been used for analysis of all data sets shown in the main text (Figure 2,3,5). For data analysis an all photon burst-search was used (M = 15, T = 500 μs and L = 50) with additional thresholding of all photons >250.

**Figure S4.**
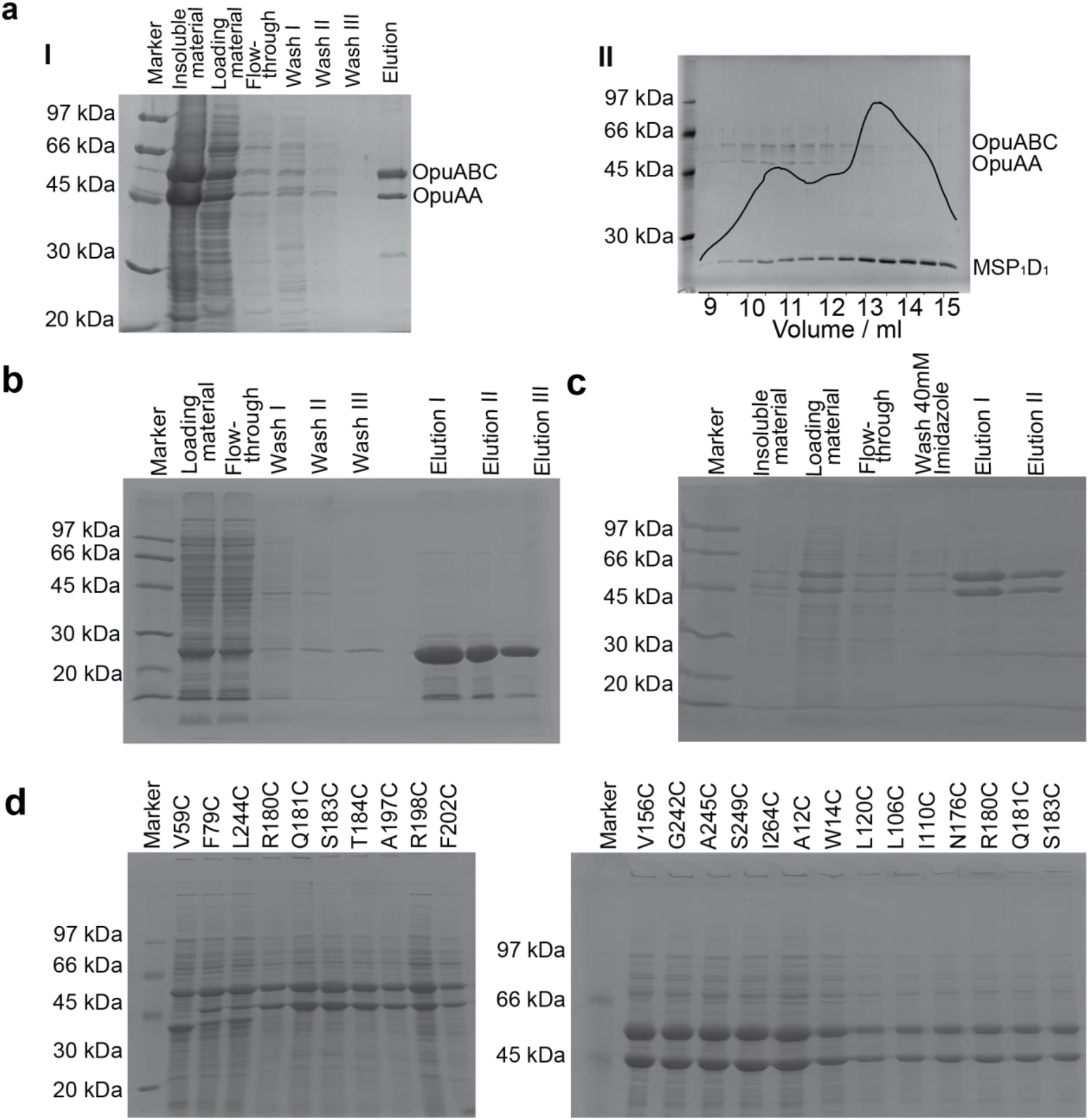
OpuA and MSP_1_D_1_ expression and purification. **(a)** SDS-PAGE of the purification process from membrane vesicles to detergent solubilized OpuA WT (I). SDS-PAGE analysis of the elution fractions of the reconstituted Nd-OpuA. The black line shows the size exclusion chromatogram and the correlation between SEC peaks and protein content (II). **(b)** SDS-PAGE from the purification of belt protein MSP_1_D_1_. At ~24 kDa **(c)** SDS-PAGE from purification of OpuA. **(d)** SDS-PAA gel electrophoresis of isolated membrane vesicles with overexpressed cysteine derivatives of OpuA. The mutants show high overexpression (OpuABC at 62KDa and OpuAA at 45 kDa) and in most cases equal ratio of ABC/AA subunits.

**Figure S5.**
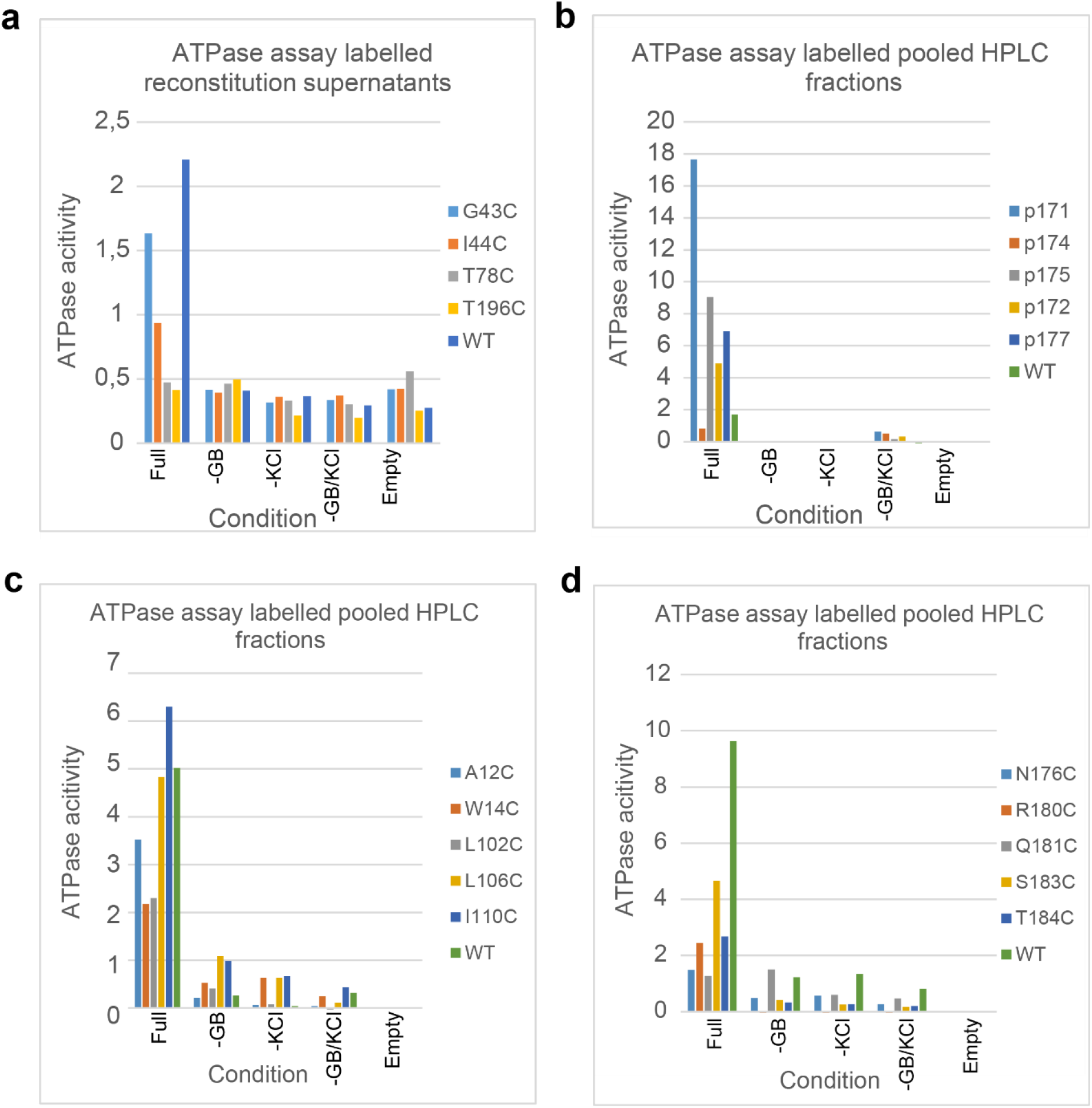
In vitro ATPase results. **(a)** This set of experiments was done without size exclusion chromatography of the reconstituted labeled nanodiscs. **(b/c/d)** These sets of assays were done with pooled fractions of nanodiscs purified with size exclusion chromatography. To determine the in vitro activity of the OpuA cysteine derivatives we used a coupled-enzyme ATPase assay (materials and methods). OpuA has been shown to be sensitive to ionic strength (4/37). To calculate the relative activity compared to WT, control reactions were done also in the absence of substrate (glycine betaine), salt (potassium chloride) and both a substrate and salt. Lastly a control with empty nanodiscs was done (only lipids and MSP_1_D_1_ present during reconstitution) to determine the baseline contribution of the nanodisc components to the assay. The ATPase assay was done as a quick way to screen through the library of mutants of valid candidates, therefore each mutant was tested once and the activity was calculated as the relative activity and not an absolute rate.

**Figure S6.**
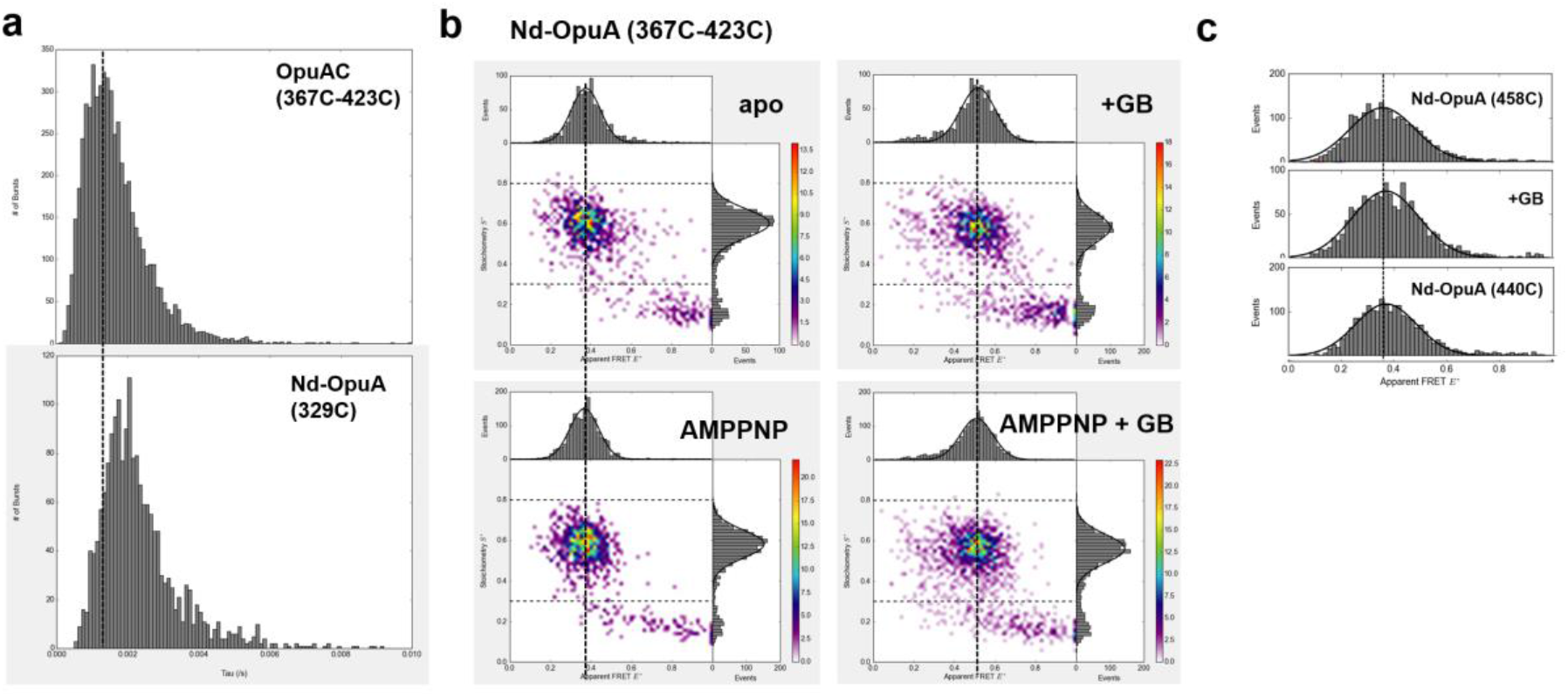
**(a)**Burst length distributions (all 100, 0.4-0.8 S) for OpuAC (free substrate protein) and Nd-OpuA (full transporter reconstituted in nanodiscs). **(b)** Additional smFRET data sets showing that AMPPNP had a similar negligible impact compared to ATP addition. **(c)** Additional smFRET data sets showing distributions of interdomain SBD labelling of Nd-OpuA (458C) and Nd-OpuA (440C).

### Supplementary Note: Homology model

The lack of a crystal structure for OpuA (TMD/NBD) led us to the use of homology modeling (Figure S7) as a tool to predict the position of amino acids in three-dimensional space. For successful fluorophore labeling, information such as solvent accessibility, steric hindrance and avoidance of proximity to the translocation channel were critical. Four structures were selected to serve as templates (Table S1) based on their transporter family, functionality, availability and sequence identity/similarity.

For each template an OpuA structure was predicted and all resulting models were quite similar with minor differences in loops connecting the trans-membrane helices (Figure S7). To assess the distance between target residues for mutagenesis and fluorophore labelling, we used the maltose transporter as template, because we had information on both inward (PDB: 2r6g) and outward (PDB: 3fh6) facing conformations. The transmembrane domains of the maltose transporter showed little variation in the inward and outward conformation (Figure 7) and were behaving as rigid domains. The produced inward and outward facing models were the basis for calculating the distance change between inward and outward conformations (Figure 7c/d). The relative distance change is critical information to maximize the resolution of the smFRET experiments.

The homology model we produced here is sensitive to the input template structures and with more template structures of high sequence identity it would produce more reliable results[1, 2]. The lack of solved transporter structures related to OpuA was thus limiting the accuracy/correctness of the produced model. To increase the likelihood of obtaining a functional mutant with good labelling behavior, the homology model was complemented with information on the conservation scores for all the amino acids in OpuA. This limited reliability led to consider an approach where we increase the number of the cysteine mutants in our library for testing (Figure 7a).

**Figure S7.**
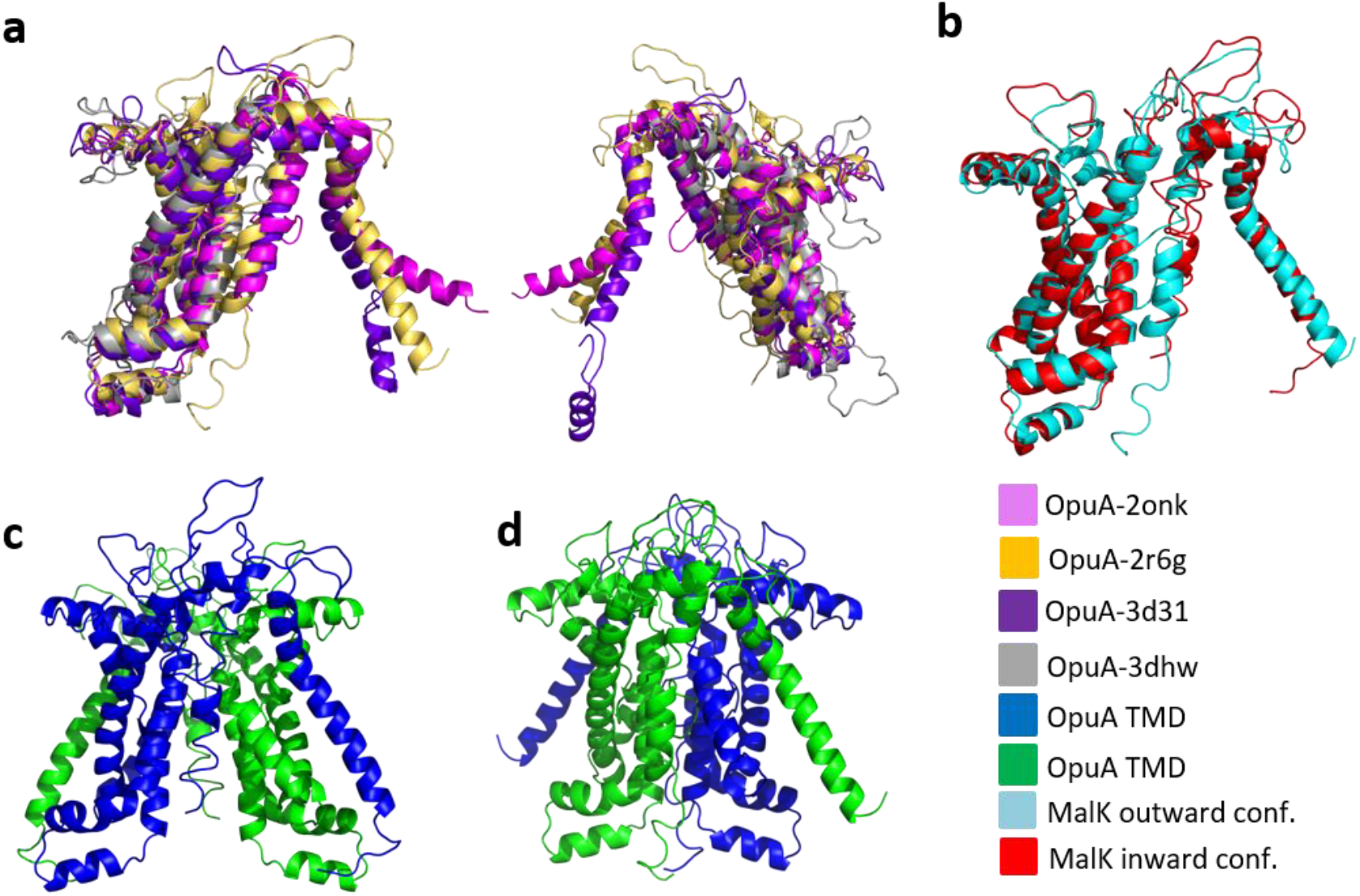
The homology model of the OpuA transmembrane domain was done using Swiss-model server [3]. The modelled structures were visualized using PyMol[4]. To further asses the quality of the model the modelled structures were aligned and overlaid on the template structures using PyMol. **(a)** Superimposition of the four modeled inward homology models based on the maltose, tungstate/molybdate, sulfate/molybdate and methionine importers. **(b)** Aligned transmembrane domains of the maltose importer in the outward and inward conformation. **(c)** Recreation of the TMD dimer in the inward facing conformation using as building blocks the X-Ray structure of the maltose-based homology model of OpuA. The orientation of the dimer is with the upper plane facing the periplasm and the lower plain facing the cytoplasm. **(d)** Recreation of the TMD dimer in the outward facing conformation based on the outward facing conformation of the maltose transporter.

**Figure S8.**
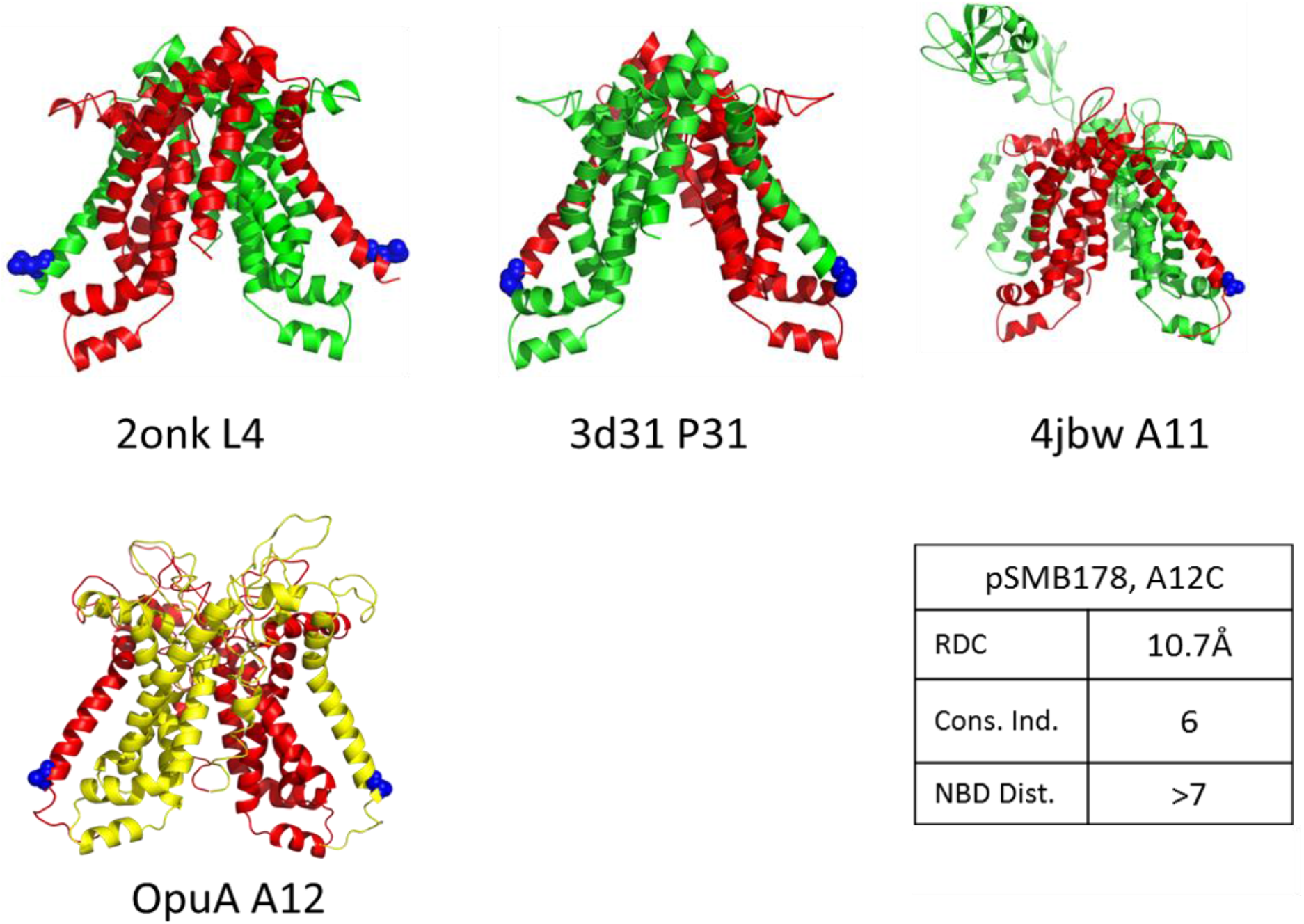
OpuA homology models with selected labelling positions relevant to monitoring TMD movements. OpuA A12C modelled inward facing dimer (OpuA A12) in red/yellow. The rest are the structures of the three templates (2onk Molybdate/tungstate ABC transporter, 3d31 Sulfate/molybdate ABC transporter, 4jbw MalG). With blue spheres are denoted the homologous amino acids to the Alanine 12 of OpuA as determined from multiple amino acid sequence alignment using ClustalW (the four template amino acid sequences were aligned with the sequence of OpuA). All the cartoons are oriented the same way, with the periplasmic space being above the cartoon and the cytoplasmic below. The cell membrane is perpendicular to the plane of the screen/paper. All the cartoons are in the inward facing conformation. RDC is realtive distance change between tha inear and outward facing conformation for Ala12. Conservation index is the conseravation score of Ala12 as calculated by ConSurf (1 being least conserved, 10 being most conserved. NBD distance is the expected distance of the Ala12 form the nucleotide binding domain based on the model (close proximity to the NBD could impair activity).

**Figure S9:**
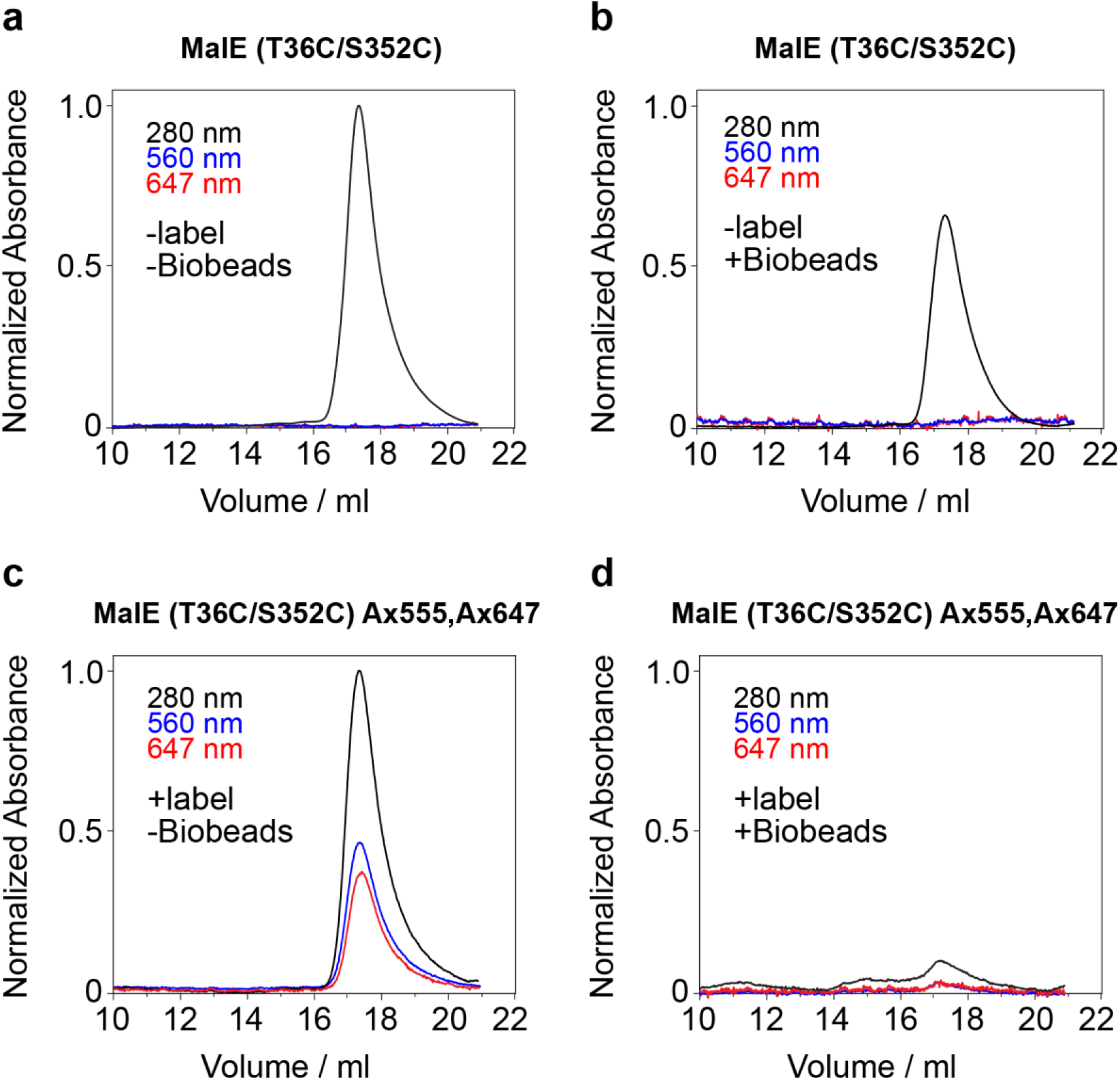
a. Chromatogram of unlabeled MalE. b. Chromatogram of unlabeled MalE after overnight incubation with SM2 bio-beads. ~34% of the total protein content was not retrieved after eluting MalE. The absorbance has been normalized against the total amount of protein of panel a. c. Labeled MalE with Ax555 and Ax647, ~78% of the available cysteines are occupied by a fluorophore. d. Labeled MalE (Ax555 and Ax647) after 3-hour incubation in SM2 bio-beads. The absorbance has been normalized against the total protein concentration of panel c.

**Table S1:** List of maleimide fluorophores used in this study including their extinction coefficients. Sources were company web-portals of Thermo-Fischer for AlexaFluor dyes[12], ATTOTEC[13], Lumiprobe[14] and ref. [15] for Cy3B.

| Fluorophore | Extinction Coefficient |
| --- | --- |
| AlexaFluor 555<sup>TM</sup> | 155000 cm<sup>-1</sup>M<sup>-1</sup> |
| AlexaFluor 647<sup>TM</sup> | 270000 cm<sup>-1</sup>M<sup>-1</sup> |
| TMR | 60000 cm<sup>-1</sup>M<sup>-1</sup> |
| Cy5 | 250000 cm<sup>-1</sup>M<sup>-1</sup> |
| Atto647N<sup>TM</sup> | 150000 cm<sup>-1</sup>M<sup>-1</sup> |
| Cy3B | 130000 cm<sup>-1</sup>M<sup>-1</sup> |

**Table S2.**
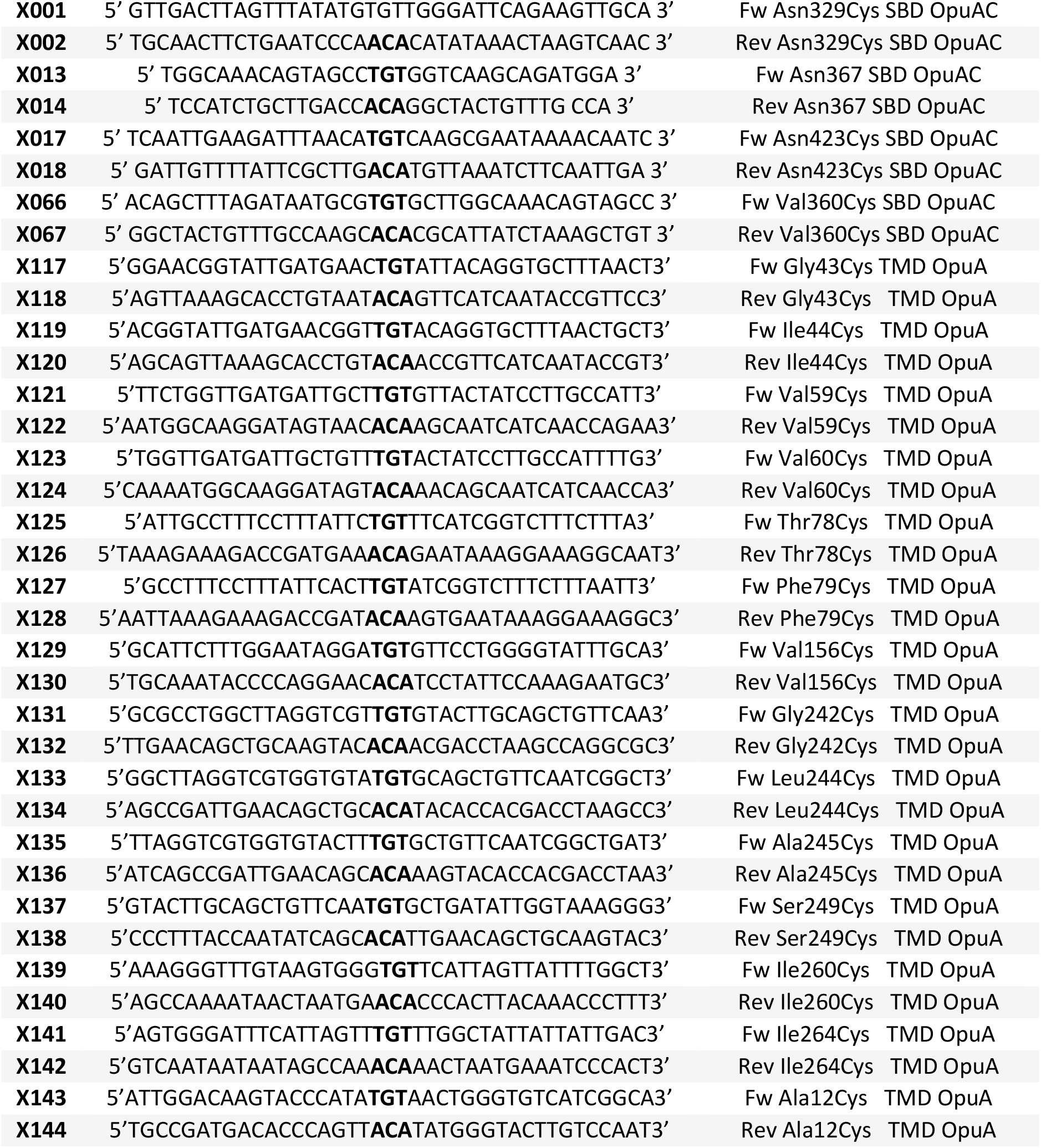
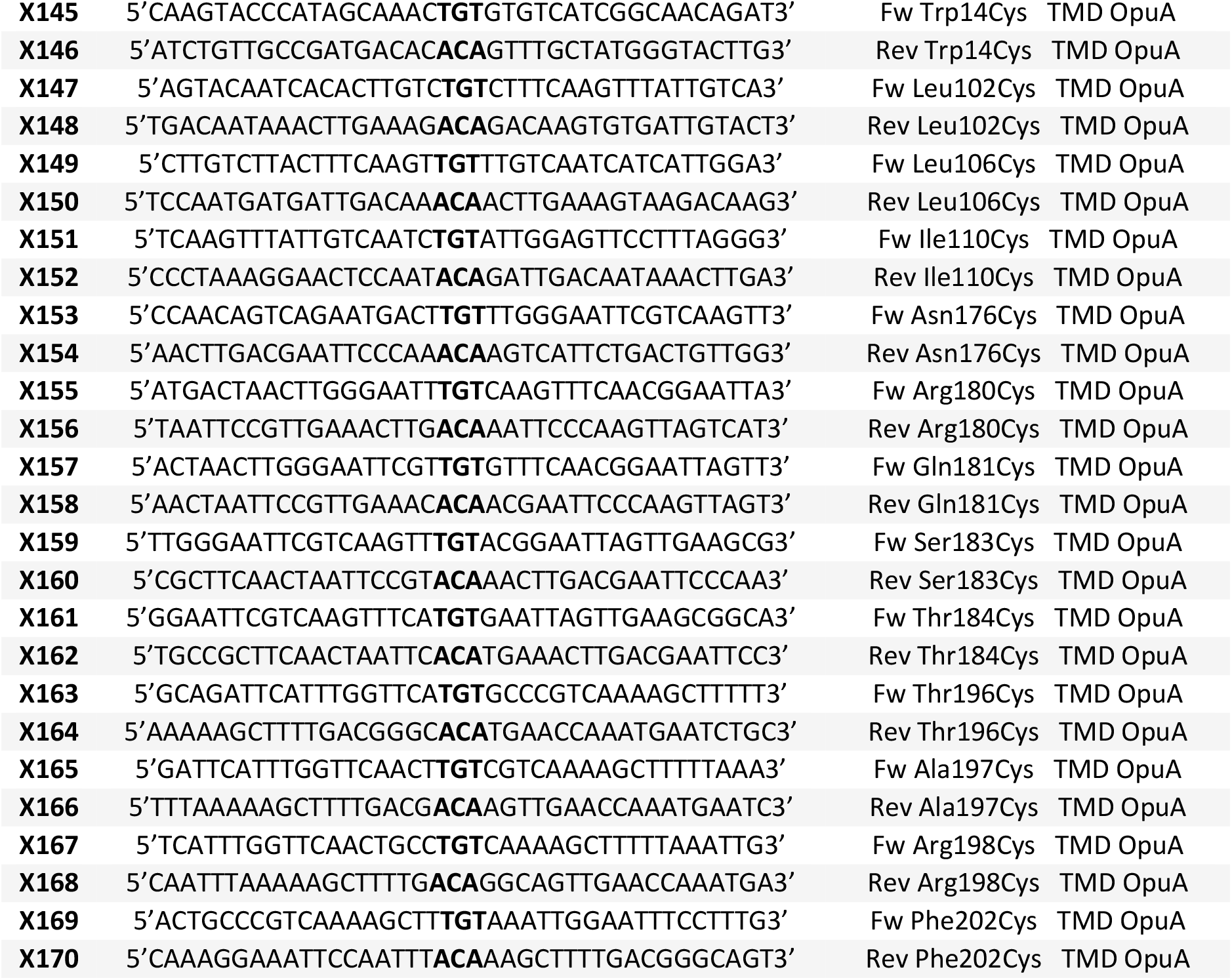
QC-PCR primers for cysteine derivatives of OpuA

**Table S3.** List of crystal structures used as templates for OpuA-TMD. The sequence identity and similarity were calculated using protein Blast. The sequence coverage score was given by swiss model and represents the percentage of the template structure that was used for the prediction of the modeled OpuA TMD. Multiple sequence alignment was done using ClustalΩ[5].

| Template | Sequence identity | Sequence similarity | Coverage | Description |
| --- | --- | --- | --- | --- |
| 2onk.1.C | 24% | 30% | 81% | Molybdate/tungstate ABC transporter, permease protein |
| 2r6g.1.E | 20% | 28% | 92% | Maltose transport system permease protein MalG (inward) |
| 4jbw.2.B | 20% | 28% | 92% | Maltose transport system permease protein MalG (outward) |
| 3d31.1.C | 26% | 29% | 84% | Sulfate/molybdate ABC transporter, permease protein |
| 3dhw.2.A | 23% | 32% | 68% | D-methionine transport system permease protein metI |

**Table S4.** Calculated distances between the cysteine positions in the homodimer of OpuA. The first distance is the relative distance change between the modeled inward- and outward-facing conformation. The second value represents the lower distance measured in either conformation. The maximum distance between the two fluorophores can be calculated by adding the relative distance change to the listed (minimum) distance. Conservation denotes the conservation index for each residue as calculated through ConSurf[6–11]. Due to the resolution range of smFRET low initial distances (less than 20Å) cannot be easily measured. However, they open the possibility of quenching assays as they are more sensitive for probes at close proximity.

| Periplasmic |  |  | Cytoplasmic |  |  |
| --- | --- | --- | --- | --- | --- |
| Mutant | net dif. / low dist. Å | Conservation score | Mutant | net dif. / low dist. Å | Conservation score |
| Gly 43 Cys | 6.4 / 48.4 outw | 1 | Ala 12 | 10.7 / 61.8 outw | 6 |
| Ile 44 Cys | 14.2 / 43.9 outw | 1 | Try 14 | 10.0 / 54.7 outw | 2 |
| Val 59 Cys | NA / NA | 1 | Leu 102 | 8.4 / 24.5 outw | 4 |
| Val 60 Cys | NA / NA | 1 | Leu 106 | 10.5 / 25.9 outw | 1 |
| Thr 78 Cys | 7.5 / 38.9 inw | 1 | Ile 110 | 8.3 / 27.8 outw | 1 |
| Phe 79 Cys | 6.1 / 39.1 inw | 5 | Asn 176 | 8.3 / 14.7 outw | 7 |
| Val 156 Cys | 6.7 / 35.8 inw | 7 | Arg 180 | 10.0 / 13.0 outw | 7 |
| Gly 242 Cys | 5.4 / 38.7 inw | 5 | Gln 181 | 9.7 / 9.2 outw | 8 |
| Leu 244 Cys | 14.1 / 29.4 inw | 8 | Ser 183 | 14.4 / 9.6 outw | 7 |
| Ala 245 Cys | 7.9 / 29.7 inw | 6 | Thr 184 | 12.4 / 15.9 outw | 1 |
| Ser 249 Cys | 5.9 / 37.2 inw | 7 | Thr 196 | 12.3 / 44.3 outw | 7 |
| Ile 260 Cys | 8.7 / 22.6 inw | 3 | Ala 197 | 13.1 / 42.3 outw | 2 |
| Ile 264 Cys | 10.6 / 15.6 outw | 7 | Arg 198 | 128 / 41.0 outw | 1 |
| Phe 202 | 13.7 / 33.6 outw | 1 |  |  |  |

## Notes

### Competing Interest Statement

The authors have declared no competing interest.

## REFERENCES

1. Schuurman-Wolters, G. K. & Poolman, B. (2005) Substrate specificity and ionic regulation of GlnPQ from Lactococcus lactis. An ATP-binding cassette transporter with four extracytoplasmic substrate-binding domains, Biol Chem. 280, 23785–90.

2. Sharom, F. J. (2008) ABC multidrug transporters: structure, function and role in chemoresistance, Pharmacogenomics. 9, 105–27.

3. Abele, R. & Tampe, R. (2004) The ABCs of immunology: structure and function of TAP, the transporter associated with antigen processing, Physiology (Bethesda). 19, 216–24.

4. Biemans-Oldehinkel, E., Doeven, M. K. & Poolman, B. (2006) ABC transporter architecture and regulatory roles of accessory domains, FEBS Lett. 580, 1023–35.

5. Davidson, A. L., Dassa, E., Orelle, C. & Chen, J. (2008) Structure, function, and evolution of bacterial ATP-binding cassette systems, Microbiol Mol Biol Rev. 72, 317–64, table of contents.

6. Oldham, M. L., Davidson, A. L. & Chen, J. (2008) Structural insights into ABC transporter mechanism, Current opinion in structural biology. 18, 726–733.

7. Kos, V. & Ford, R. C. (2009) The ATP-binding cassette family: a structural perspective, Cell Mol Life Sci. 66, 3111–26.

8. Parcej, D. & Tampe, R. (2010) ABC proteins in antigen translocation and viral inhibition, Nat Chem Biol. 6, 572–80.

9. Locher, K. P. (2009) Review. Structure and mechanism of ATP-binding cassette transporters, Philos Trans R Soc Lond B Biol Sci. 364, 239–45.

10. Plumptre, C. D., Eijkelkamp, B. A., Morey, J. R., Behr, F., Couñago, R. M., Ogunniyi, A. D., Kobe, B., O’Mara, M. L., Paton, J. C. & McDevitt, C. A. (2014) AdcA and AdcAII employ distinct zinc acquisition mechanisms and contribute additively to zinc homeostasis in S treptococcus pneumoniae, Molecular microbiology. 91, 834–851.

11. Riordan, J. R. (2008) CFTR function and prospects for therapy, Annu Rev Biochem. 77, 701–26.

12. Teague, S. J. (2003) Implications of protein flexibility for drug discovery, Nature reviews Drug discovery. 2, 527–541.

13. Majumder, P., Mallela, A. K. & Penmatsa, A. (2018) Transporters Through the Looking Glass: An Insight into the Mechanisms of Ion-Coupled Transport and Methods That Help Reveal Them, Journal of the Indian Institute of Science. 98, 283–300.

14. Majumdar, D. S., Smirnova, I., Kasho, V., Nir, E., Kong, X., Weiss, S. & Kaback, H. R. (2007) Singlemolecule FRET reveals sugar-induced conformational dynamics in LacY, Proc Natl Acad Sci U S A. 104, 12640–5.

15. Zhao, Y., Terry, D., Shi, L., Weinstein, H., Blanchard, S. C. & Javitch, J. A. (2010) Single-molecule dynamics of gating in a neurotransmitter transporter homologue, Nature. 465, 188–93.

16. Zhao, Y., Terry, D. S., Shi, L., Quick, M., Weinstein, H., Blanchard, S. C. & Javitch, J. A. (2011) Substrate-modulated gating dynamics in a Na+-coupled neurotransmitter transporter homologue, Nature. 474, 109–13.

17. Akyuz, N., Altman, R. B., Blanchard, S. C. & Boudker, O. (2013) Transport dynamics in a glutamate transporter homologue, Nature. 502, 114–8.

18. Erkens, G. B., Hanelt, I., Goudsmits, J. M., Slotboom, D. J. & van Oijen, A. M. (2013) Unsynchronised subunit motion in single trimeric sodium-coupled aspartate transporters, Nature. 502, 119–23.

19. Gouridis, G., Schuurman-Wolters, G. K., Ploetz, E., Husada, F., Vietrov, R., de Boer, M., Cordes, T. & Poolman, B. (2015) Conformational dynamics in substrate-binding domains influences transport in the ABC importer GlnPQ, Nat Struct Mol Biol. 22, 57–64.

20. van der Velde, J. H., Oelerich, J., Huang, J., Smit, J. H., Aminian Jazi, A., Galiani, S., Kolmakov, K., Guoridis, G., Eggeling, C., Herrmann, A., Roelfes, G. & Cordes, T. (2016) A simple and versatile design concept for fluorophore derivatives with intramolecular photostabilization, Nat Commun. 7, 10144.

21. de Boer, M., Gouridis, G., Vietrov, R., Begg, S. L., Schuurman-Wolters, G. K., Husada, F., Eleftheriadis, N., Poolman, B., McDevitt, C. A. & Cordes, T. (2019) Conformational and dynamic plasticity in substrate-binding proteins underlies selective transport in ABC importers, Elife. 8, e44652.

22. Jazi, A. A., Ploetz, E., Arizki, M., Dhandayuthapani, B., Waclawska, I., Krämer, R., Ziegler, C. & Cordes, T. (2017) Caging and Photoactivation in Single-Molecule Förster Resonance Energy Transfer Experiments, Biochemistry. 56, 2031–2041.

23. Husada, F., Bountra, K., Tassis, K., de Boer, M., Romano, M., Rebuffat, S., Beis, K. & Cordes, T. (2018) Conformational dynamics of the ABC transporter McjD seen by single-molecule FRET, The EMBO journal. 37, e100056.

24. Yang, M., Livnat Levanon, N., Acar, B., Aykac Fas, B., Masrati, G., Rose, J., Ben-Tal, N., Haliloglu, T., Zhao, Y. & Lewinson, O. (2018) Single-molecule probing of the conformational homogeneity of the ABC transporter BtuCD, Nat Chem Biol. 14, 715–722.

25. Liu, Y., Liu, Y., He, L., Zhao, Y. & Zhang, X. C. (2018) Single-molecule fluorescence studies on the conformational change of the ABC transporter MsbA, Biophysics Reports. 4, 153–165.

26. Ha, T., Enderle, T., Ogletree, D., Chemla, D. S., Selvin, P. R. & Weiss, S. (1996) Probing the interaction between two single molecules: fluorescence resonance energy transfer between a single donor and a single acceptor, Proceedings of the National Academy of Sciences. 93, 6264–6268.

27. Ha, T. (2001) Single-Molecule FRET, Single Molecules. 2, 283–284.

28. Lerner, E., Cordes, T., Ingargiola, A., Alhadid, Y., Chung, S., Michalet, X. & Weiss, S. (2018) Toward dynamic structural biology: Two decades of single-molecule Förster resonance energy transfer, Science. 359.

29. Muschielok, A., Andrecka, J., Jawhari, A., Bruckner, F., Cramer, P. & Michaelis, J. (2008) A nanopositioning system for macromolecular structural analysis, Nat Methods. 5, 965–71.

30. Kalinin, S., Peulen, T., Sindbert, S., Rothwell, P. J., Berger, S., Restle, T., Goody, R. S., Gohlke, H. & Seidel, C. A. (2012) A toolkit and benchmark study for FRET-restrained high-precision structural modeling, Nat Methods. 9, 1218–25.

31. Hellenkamp, B., Schmid, S., Doroshenko, O., Opanasyuk, O., Kühnemuth, R., Adariani, S. R., Ambrose, B., Aznauryan, M., Barth, A. & Birkedal, V. (2018) Precision and accuracy of single-molecule FRET measurements—a multi-laboratory benchmark study, Nature methods. 15, 669–676.

32. Cordes, T., Santoso, Y., Tomescu, A. I., Gryte, K., Hwang, L. C., Camará, B., Wigneshweraraj, S. & Kapanidis, A. N. (2010) Sensing DNA opening in transcription using quenchable Forster resonance energy transfer, Biochemistry. 49, 9171–9180.

33. Kapanidis, A. N., Lee, N. K., Laurence, T. A., Doose, S., Margeat, E. & Weiss, S. (2004) Fluorescence-aided molecule sorting: analysis of structure and interactions by alternating-laser excitation of single molecules, Proc Natl Acad Sci U S A. 101, 8936–41.

34. Robb, N. C., Cordes, T., Hwang, L. C., Gryte, K., Duchi, D., Craggs, T. D., Santoso, Y., Weiss, S., Ebright, R. H. & Kapanidis, A. N. (2013) The transcription bubble of the RNA polymerase-promoter open complex exhibits conformational heterogeneity and millisecond-scale dynamics: implications for transcription start-site selection, Journal of molecular biology. 425, 875–885.

35. Gouridis, G., Hetzert, B., Kiosze-Becker, K., de Boer, M., Heinemann, H., Nurenberg-Goloub, E., Cordes, T. & Tampe, R. (2019) ABCE1 Controls Ribosome Recycling by an Asymmetric Dynamic Conformational Equilibrium, Cell Rep. 28, 723–734 e6.

36. Goudsmits, J. M., Slotboom, D. J. & van Oijen, A. M. (2017) Single-molecule visualization of conformational changes and substrate transport in the vitamin B 12 ABC importer BtuCD-F, Nature communications. 8, 1–10.

37. Ruan, Y., Miyagi, A., Wang, X., Chami, M., Boudker, O. & Scheuring, S. (2017) Direct visualization of glutamate transporter elevator mechanism by high-speed AFM, Proc Natl Acad Sci U S A. 114, 1584–1588.

38. Joseph, B., Sikora, A., Bordignon, E., Jeschke, G., Cafiso, D. S. & Prisner, T. F. (2015) Distance measurement on an endogenous membrane transporter in E. coli cells and native membranes using EPR spectroscopy, Angewandte Chemie. 127, 6294–6297.

39. Sahu, I. D., McCarrick, R. M., Troxel, K. R., Zhang, R., Smith, H. J., Dunagan, M. M., Swartz, M. S., Rajan, P. V., Kroncke, B. M. & Sanders, C. R. (2013) DEER EPR measurements for membrane protein structures via bifunctional spin labels and lipodisq nanoparticles, Biochemistry. 52, 6627–6632.

40. Karasawa, A., Swier, L. J., Stuart, M. C., Brouwers, J., Helms, B. & Poolman, B. (2013) Physicochemical factors controlling the activity and energy coupling of an ionic strength-gated ATP-binding cassette (ABC) transporter, Biol Chem. 288, 29862–71.

41. Biemans-Oldehinkel, E., Mahmood, N. A. & Poolman, B. (2006) A sensor for intracellular ionic strength, Proc Natl Acad Sci U S A. 103, 10624–9.

42. Patzlaff, J. S., van der Heide, T. & Poolman, B. (2003) The ATP/substrate stoichiometry of the ATP-binding cassette (ABC) transporter OpuA, Biol Chem. 278, 29546–51.

43. Heide, T., Stuart, M. C. & Poolman, B. (2001) On the osmotic signal and osmosensing mechanism of an ABC transport system for glycine betaine, The EMBO Journal. 20, 7022–32.

44. van der Fulyani, F., Schuurman-Wolters, G. K., Slotboom, D.-J. & Poolman, B. (2016) Relative rates of amino acid Import via the ABC transporter GlnPQ determine the growth performance of Lactococcus lactis, Journal of bacteriology. 198, 477–485.

45. Nguyen, P. T., Lai, J. Y., Lee, A. T., Kaiser, J. T. & Rees, D. C. (2018) Noncanonical role for the binding protein in substrate uptake by the MetNI methionine ATP Binding Cassette (ABC) transporter, Proceedings of the National Academy of Sciences. 115, E10596–E10604.

46. Karasawa, A., Erkens, G. B., Berntsson, R. P., Otten, R., Schuurman-Wolters, G. K., Mulder, F. A. & Poolman, B. (2011) Cystathionine beta-synthase (CBS) domains 1 and 2 fulfill different roles in ionic strength sensing of the ATP-binding cassette (ABC) transporter OpuA, Biol Chem. 286, 37280–91.

47. Kijac, A., Shih, A. Y., Nieuwkoop, A. J., Schulten, K., Sligar, S. G. & Rienstra, C. M. (2010) Lipid-protein correlations in nanoscale phospholipid bilayers determined by solid-state nuclear magnetic resonance, Biochemistry. 49, 9190–9198.

48. ter Beek, J., Guskov, A. & Slotboom, D. J. (2014) Structural diversity of ABC transporters, Gen Physiol. 143, 419–35.

49. Le Reste, L., Hohlbein, J., Gryte, K. & Kapanidis, A. N. (2012) Characterization of dark quencher chromophores as nonfluorescent acceptors for single-molecule FRET, Biophysical journal. 102, 2658–2668.

50. Ploetz, E., Lerner, E., Husada, F., Roelfs, M., Chung, S., Hohlbein, J., Weiss, S. & Cordes, T. (2016) Förster resonance energy transfer and protein-induced fluorescence enhancement as synergetic multi-scale molecular rulers, Scientific reports. 6, 33257.

51. Bertoni, M., Kiefer, F., Biasini, M., Bordoli, L. & Schwede, T. (2017) Modeling protein quaternary structure of homo-and hetero-oligomers beyond binary interactions by homology, Scientific reports. 7, 1–15.

52. Guex, N., Peitsch, M. C. & Schwede, T. (2009) Automated comparative protein structure modeling with SWISS-MODEL and Swiss-PdbViewer: A historical perspective, Electrophoresis. 30, S162–S173.

53. Häusler, E., Fredriksson, K., Goba, I., Peters, C., Raltchev, K., Sperl, L., Steiner, A., Weinkauf, S. & Hagn, F. (2020) Quantifying the insertion of membrane proteins into lipid bilayer nanodiscs using a fusion protein strategy, Biochimica et Biophysica Acta-Biomembranes. 1862, 183190.

54. Lewinson, O. & Livnat-Levanon, N. (2017) Mechanism of action of ABC importers: conservation, divergence, and physiological adaptations, Journal of molecular biology. 429, 606–619.

55. Szöllősi, D., Rose-Sperling, D., Hellmich, U. A. & Stockner, T. (2018) Comparison of mechanistic transport cycle models of ABC exporters, Biochimica et Biophysica Acta-Biomembranes. 1860, 818–832.

56. Scheepers, G. H., Lycklama a Nijeholt, J. A. & Poolman, B. (2016) An updated structural classification of substrate-binding proteins, FEBS letters. 590, 4393–4401.

57. Davidson, A. L., Shuman, H. A. & Nikaido, H. (1992) Mechanism of maltose transport in Escherichia coli: transmembrane signaling by periplasmic binding proteins, Proceedings of the National Academy of Sciences. 89, 2360–2364.

58. Borths, E. L., Poolman, B., Hvorup, R. N., Locher, K. P. & Rees, D. C. (2005) In vitro functional characterization of BtuCD-F, the Escherichia coli ABC transporter for vitamin B12 uptake, Biochemistry. 44, 16301–16309.

59. Vigonsky, E., Ovcharenko, E. & Lewinson, O. (2013) Two molybdate/tungstate ABC transporters that interact very differently with their substrate binding proteins, Proceedings of the National Academy of Sciences. 110, 5440–5445.

60. Gul, N., Schuurman-Wolters, G., Karasawa, A. & Poolman, B. (2012) Functional characterization of amphipathic α-helix in the osmoregulatory ABC transporter OpuA, Biochemistry. 51, 5142–5152.

61. Spack Jr, E. G., Packard, B., Wier, M. L. & Edidin, M. (1986) Hydrophobic adsorption chromatography to reduce nonspecific staining by rhodamine-labeled antibodies, Analytical biochemistry. 158, 233–237.

62. DeLano, W. L. (2002) Pymol: An open-source molecular graphics tool, Newsletter on protein crystallography. 40, 82–92.

63. de Boer, M., Gouridis, G., Muthahari, Y. A. & Cordes, T. (2019) Single-molecule observation of ligand binding and conformational changes in FeuA, Biophysical journal. 117, 1642–1654.

64. van der Velde, J. H., Ploetz, E., Hiermaier, M., Oelerich, J., de Vries, J. W., Roelfes, G. & Cordes, T. (2013) Mechanism of intramolecular photostabilization in self-healing cyanine fluorophores, ChemPhysChem. 14, 4084–4093.

65. Rabiner, L. R. (1990) A tutorial on hidden markov models and selected applications in speech recognition. Readings in Speech Recognition (pp. 267–296) in, Los Altos, CA: Morgan Kaufmann,

## References

1. Bordoli, L., Kiefer, F., Arnold, K., Benkert, P., Battey, J. & Schwede, T. (2009) Protein structure homology modeling using SWISS-MODEL workspace, Nature protocols. 4, 1.

2. Schwede, T., Kopp, J., Guex, N. & Peitsch, M. C. (2003) SWISS-MODEL: an automated protein homology-modeling server, J Nucleic acids research. 31, 3381–3385.

3. Guex, N., Peitsch, M. C. & Schwede, T. (2009) Automated comparative protein structure modeling with SWISS-MODEL and Swiss-PdbViewer: A historical perspective, J Electrophoresis. 30, S162–S173.

4. Delano, W. L. (2002) The PyMOL molecular graphics system, version 1.8, Schrödinger LLC. 10.

5. Sievers, F., Wilm, A., Dineen, D., Gibson, T. J., Karplus, K., Li, W., Lopez, R., McWilliam, H., Remmert, M. & Söding, J. (2011) Fast, scalable generation of high-quality protein multiple sequence alignments using Clustal Omega, J Molecular systems biology. 7, 539.

6. Ashkenazy, H., Abadi, S., Martz, E., Chay, O., Mayrose, I., Pupko, T. & Ben-Tal, N. (2016) ConSurf 2016: an improved methodology to estimate and visualize evolutionary conservation in macromolecules, Nucleic acids research. 44, W344–W350.

7. Ashkenazy, H., Erez, E., Martz, E., Pupko, T. & Ben-Tal, N. (2010) ConSurf 2010: calculating evolutionary conservation in sequence and structure of proteins and nucleic acids, Nucleic acids research. 38, W529–W533.

8. Berezin, C., Glaser, F., Rosenberg, J., Paz, I., Pupko, T., Fariselli, P., Casadio, R. & Ben-Tal, N. (2004) ConSeq: the identification of functionally and structurally important residues in protein sequences, J Bioinformatics. 20, 1322–1324.

9. Celniker, G., Nimrod, G., Ashkenazy, H., Glaser, F., Martz, E., Mayrose, I., Pupko, T. & Ben-Tal, N. (2013) ConSurf: using evolutionary data to raise testable hypotheses about protein function, Israel Journal of Chemistry. 53, 199–206.

10. Glaser, F., Pupko, T., Paz, I., Bell, R. E., Bechor-Shental, D., Martz, E. & Ben-Tal, N. (2003) ConSurf: identification of functional regions in proteins by surface-mapping of phylogenetic information, J Bioinformatics. 19, 163–164.

11. Landau, M., Mayrose, I., Rosenberg, Y., Glaser, F., Martz, E., Pupko, T. & Ben-Tal, N. (2005) ConSurf 2005: the projection of evolutionary conservation scores of residues on protein structures, J Nucleic acids research. 33, W299–W302.

12. https://www.thermofisher.com/de/de/home/references/molecular-probes-the-handbook/technical-notes-and-product-highlights/the-alexa-fluor-dye-series.html

13. https://www.atto-tec.com/product_info.php?info=p114_atto-647n.html

14. https://de.lumiprobe.com/fluorophore-chart

15. Cooper, M., Ebner, A., Briggs, M., Burrows, M., Gardner, N., Richardson, R. & West, R. (2004) Cy3B^TM^: Improving the Performance of Cyanine Dyes. Journal of Fluorescence, 14, 145–150.

